# PRISM: Recovering cell type specific expression profiles from composite RNA-seq data

**DOI:** 10.1101/854505

**Authors:** Antti Häkkinen, Kaiyang Zhang, Amjad Alkodsi, Noora Andersson, Erdogan Pekcan Erkan, Jun Dai, Katja Kaipio, Tarja Lamminen, Naziha Mansuri, Kaisa Huhtinen, Anna Vähärautio, Olli Carpén, Johanna Hynninen, Sakari Hietanen, Rainer Lehtonen, Sampsa Hautaniemi

## Abstract

A major challenge in analyzing cancer patient transcriptomes is that the tumors are inherently heterogeneous and evolving. We analyzed 214 bulk RNA samples of a longitudinal, prospective ovarian cancer cohort and found that the sample composition changes systematically due to chemotherapy and between the anatomical sites, preventing direct comparison of treatment-naive and treated samples. To overcome this, we developed PRISM, a latent statistical framework to simultaneously extract the sample composition and cell type specific whole-transcriptome profiles adapted to each individual sample. Our results indicate that the PRISM-derived composition-free transcriptomic profiles and signatures derived from them predict the patient response better than the composite raw bulk data. We validated our findings in independent ovarian cancer and melanoma cohorts, and verified that PRISM accurately estimates the composition and cell type specific expression through whole-genome sequencing and RNA *in situ* hybridization experiments. PRISM is freely available with full source code and documentation.

---

Precision oncology aims to identify targetable alterations based on molecular profiling of tumors ^1^. As cancers are heterogeneous diseases that evolve during treatment and follow-up ^2, 3^, an essential part of precision oncology is the use of transcriptomic data from samples collected before, during, and after therapy ^4, 5^. However, a major unresolved challenge in analyzing longitudinal data is that the sample composition, i.e. the fraction of cancer, stromal, and immune cells, in the patient-derived samples varies significantly, which severely hinders subsequent analyses ^6^.

Alleviating the sample composition issue by discarding low tumor content samples ^7, 8^ can bias the sampling to contain only cancer cell rich tumors and exclude samples from good-responding patients during therapy, which is detrimental in longitudinal cohorts. Current computational correction approaches are not ideally suited for precision oncology needs as they focus on either immune or stromal signatures and employ preset expression profiles ^9–11^, derive the sample composition without estimating the transcriptomic profiles ^12, 13^, operate at a population level only ^14^, or lack ability to adapt to patients lacking a matched single-cell data ^14, 15^.

Here, we report a novel statistical framework PRISM (Poisson RNA-profile Identification in Scaled Mixtures) for addressing the sample composition issue. PRISM is freely available and is unique in that it allows extracting bulk specific expression profiles for each constituent cell type along with their composition, at an individual sample level, by exploiting a single-cell population (potentially unmatched) reference. The single-cell reference is treated as data and is also subjected to the statistical model, which combines the patterns in the single-cell data into an adaptive patient specific expression profiles.

We applied PRISM on 214 bulk RNA-seq samples that were longitudinally collected from homogeneously treated high-grade serous ovarian cancer (HGSOC) patients. HGSOC is the most common subtype of epithelial ovarian cancer with only 43% five-year survival rate ^16^. It is also one of the most genomically heterogeneous cancers ^7, 17^, characterized by high number of structural changes ^7, 17^, highlighting the importance of transcriptomic analysis and the challenges in sample comparison. Our results show that the PRISM-estimated HGSOC cell type specific expression profiles and cancer subtypes derived from them better predict disease progression than those from the composite raw bulk data. Finally, after validating the accuracy of the compositional estimates using whole-genome sequencing (WGS) data and the cell type specificity of expression levels using RNA *in situ* hybridization (RNA-ish) experiments, we confirmed that the improved survival prediction generalizes to other cohorts and cancer types by using publicly available data from The Cancer Genome Atlas (TCGA).

## Results

### PRISM: A latent statistical framework for recovering cell type specific expression profiles from RNA-seq data

PRISM employs a latent statistical model for composite (a mixture of multiple phenotypes) RNA-seq data, which accounts biological heterogeneity, compositional heterogeneity, and sampling noise. The estimated model can be exploited for decomposing bulk RNA-seq data, finding sample specific scale factors, or clustering RNA-seq data. An overview of PRISM is shown in Fig. 1, details are given in Methods, and derivation in Supplement. Briefly, given a bulk RNA sample, PRISM estimates the frequency and a sample specific whole-transcriptome profile for each cell type, by exploiting labeled a set of heterogeneous single-cell data for the desired cell types. The single-cell data need not be from matching tumors, but a set of sample capturing the between-patient heterogeneity in each cell type suffices. In the absence of labels, PRISM can derive a labeling through clustering (evaluated in Supplement). PRISM is available at https://bitbucket.org/anthakki/prism/ under the simplified BSD license.

**Figure 1.**
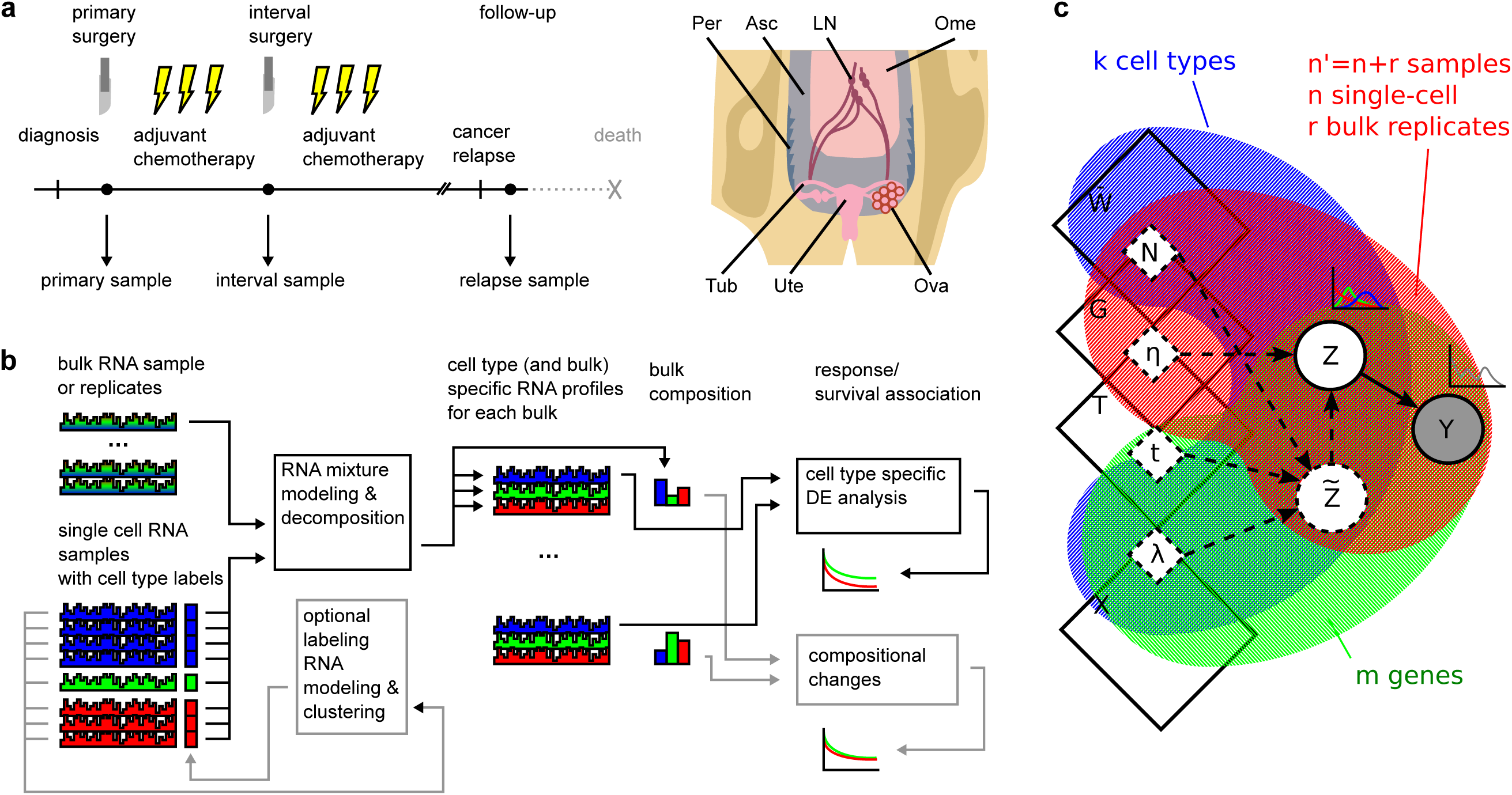
Overview of the sample collection, data analysis, and the PRISM model. a) Samples are collected from high-grade serous ovarian cancer (HGSOC) patients before neoadjuvant chemotherapy (120 samples), after three rounds of chemotherapy in the interval debulking surgery (60) and from relapsed cancers (20). For reference, we used single-cell RNA-seq data from eight matched samples (6,312 cells). Anatomical locations of the samples are indicated as follows: Asc (ascites), LN (lymph node), Ome (omentum), Ova (ovary), Per (peritoneum), Tub (fallopian tube), Ute (uterus). b) PRISM allows decomposing each bulk sample using a panel of single-cell samples, revealing the bulk compositions and expression profiles for each constituent cell type. Afterwards, differential expression or the compositional differences can be associated with patient response and survival independently. c) Plate graph for the PRISM framework described by the physical constants, *i.e.*, number of cells (*N*), sampling efficiency (*η*), expression variability (*t*^−1^), and expression mean (*λ*) generating the latent RNA count 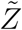 and readout *Z* for each gene and cell type in a sample. As the physical parameters are not identifiable, we parametrize the problem using mean expression (*X*), readout precision (*T*), sample scaling factor (*G*), and relative sample composition 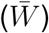. These parameters can be estimated from a set of mixture readouts (*Y*), which need not to be unimodal, by assuming the cell type specific readouts (*Z*) are scaled Poisson distributed.

### Tumor composition depends systematically on the treatment phase and the anatomical location

We first studied how the composition of HGSOC bulk samples varies over the treatment phase, the anatomical location, and the treatment response (cf. Fig. 1a). Fig. 2 shows the distribution of the PRISM-derived sample compositions. Samples taken before the treatment contain ∼70% cancer cells, while the interval samples taken after neoadjuvant chemotherapy (NACT) contain only ∼40% cancer cells, along with more fibroblasts and immune cells, and the relapse samples contain more cancer and immune cells than the treatment-naive and interval samples. This is expected, as HGSOC is typically diagnosed at advanced stage with high tumor burden, and ∼80% of the patients respond well to the first-line therapy ^18^. The results reveal, however, that a direct comparison of treatment-naive and interval samples without compositional analysis is severely biased by the compositional changes.

**Figure 2.**
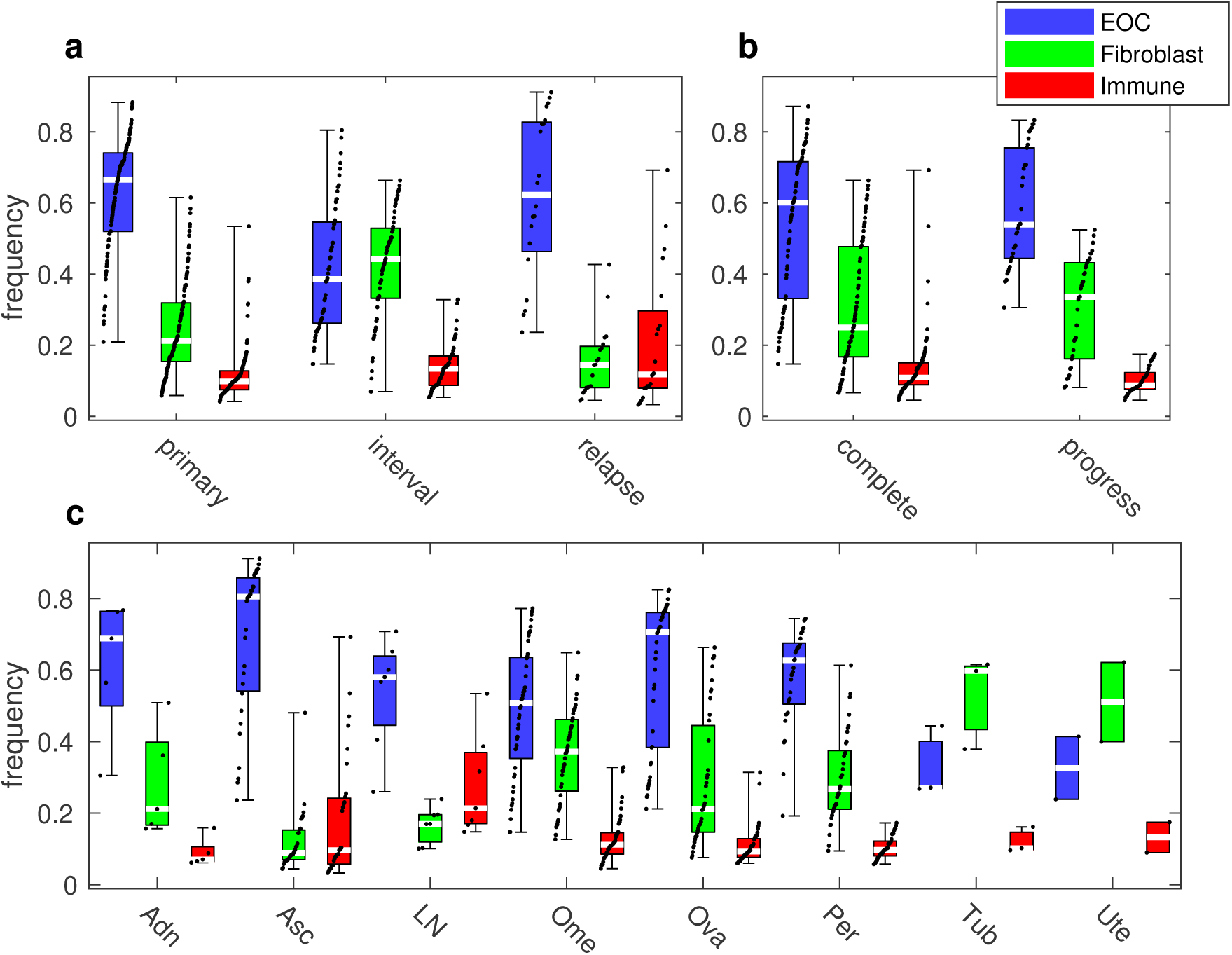
Composition of bulk RNA tissue samples in HGSOC patients. a) Cancer, fibroblast, and immune cell frequency by treatment phase: treatment-naive (primary), after three rounds of neoadjuvant chemotherapy (interval) or relapse. b) Composition by the treatment outcome: complete response (complete) or progressive disease (progress). c) Composition by the anatomical site: adnex (Adn), ascites (Asc), lymph node (LN), omentum (Ome), ovary (Ova), peritoneum (Per), fallopian tube (Tub), uterus (Ute). The boxes represent first to third quartile, white lines the medians, and whiskers the data range. Black dots represent all data, jittered by their rank.

Specifically, the fraction of cancer cells and fibroblasts vary significantly between the treatment-naive and interval samples, even when accounting for anatomical sites (p-value *p*_rc_ < 2.6 ⋅ 10^−6^ for no partial rank correlation in a t-test), whereas the number of immune cells does not (*p*_rc_ = 0.74). Similarly, we found a significant difference between the interval and relapsed cancers (*p*_rc_ < 7.1 ⋅ 10^−3^), but no difference between the primary and the relapsed samples (*p*_rc_ ∈ [0.070, 0.72]), when accounting for the anatomical site. We also quantified, for the first time, the impact of anatomical sites to the sample composition: omentum, ovary, and peritoneum have similar composition (*p*_rc_ ∈ [0.066, 0.45]), when accounting for the treatment phase differences. Also fallopian tube and uterus are similar with each other, whereas the composition of the ascites samples differs significantly from the solid samples (*p*_*rc*_ < 2.9 ⋅ 10^−4^).

Tumor composition differences between the complete response versus progressive disease groups are explained completely by the variations in the treatment phase and anatomical site of the sample (*p*_rc_ ∈ [0.092, 0.93]), which both contribute independent variation. Consequently, we argue that the composition of a patient bulk tissue sample is a strong confounder, but not a major predictive factor the patient response, necessitating expression profile analysis that controls for the sample composition.

### Decomposing bulk RNA-seq data enables cell type specific gene expression analysis

Next, we examined the PRISM-derived cell type specific expression profiles in the cancer, stromal, and immune cells. Fig. 3a shows that the expression levels of well-known cell type specific genes are higher in the respective cell type (p-value *p*_m_ < 1.9 ⋅ 10^−15^ for equal medians in a rank-sum test), and that the cell type specific expression is enriched in the decomposed profiles with respect to the composite bulk (*p*_m_ < 8.0 ⋅ 10^−4^). These imply that, the composite expression signal is also diluted by the presence of non-specific signals, masking cell type specific phenotypic changes. The cell type specificity of known housekeeping genes ^19^ is significantly lower than other genes with comparable expression level (*p*_m_ < 5.2 ⋅ 10^−6^), suggesting the specificity is well-founded.

**Figure 3.**
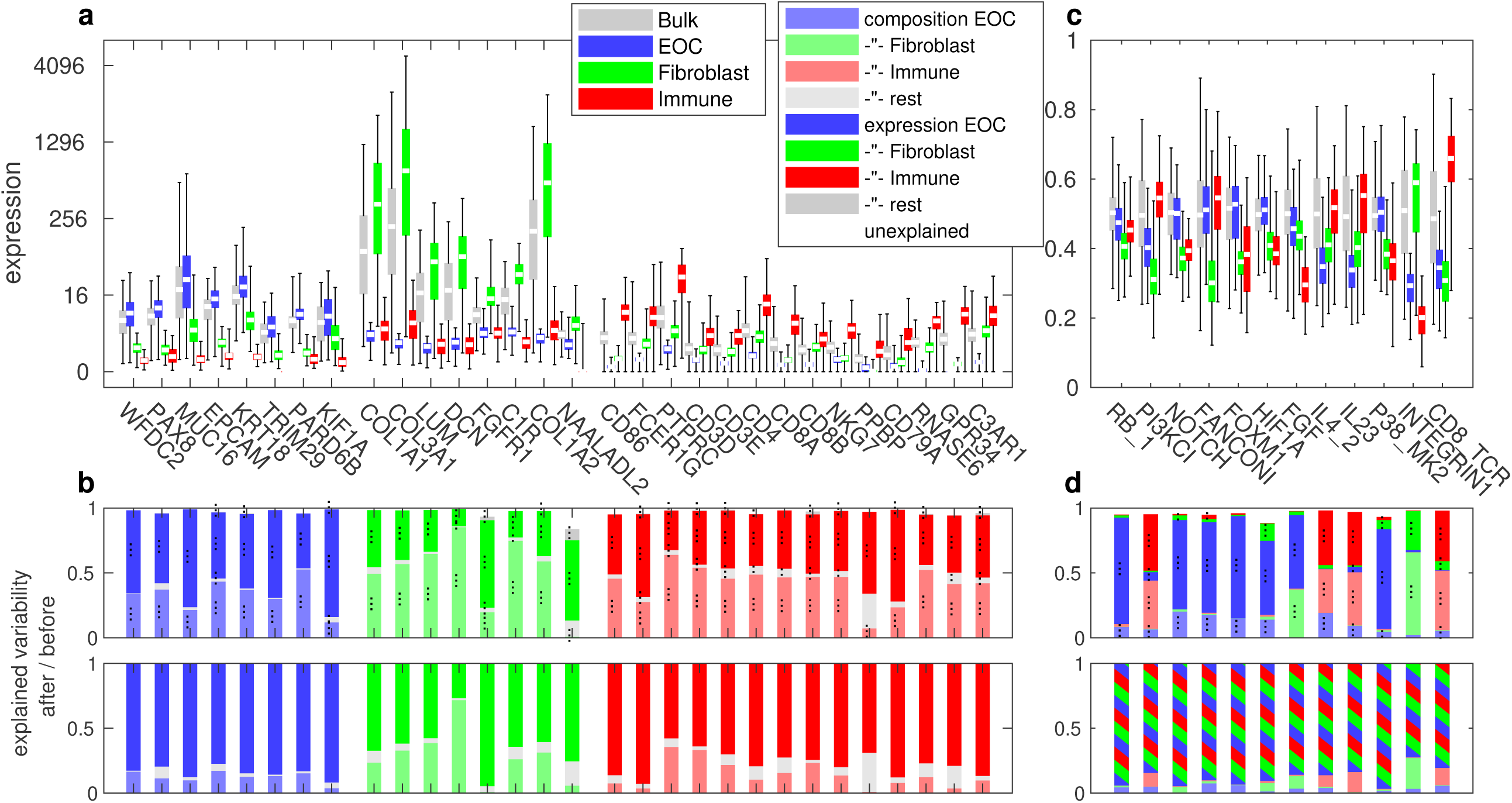
Expression in the composite bulk and the decomposed, cell type specific signals. a) Distribution of expression levels by cell type. Box is first to third quartile, white is median, whiskers are all data. b) Breakdown of the (rank) expression variability before (upper panel) or after (lower panel) the decomposition. The first group of genes represents cancer specific genes, while the second and third fibroblast and immune specific genes, respectively. c) and d) show the corresponding relative pathway activity using normalized GSEA ^20^ scores. Two or three dots indicate significance at 10^−2^ and 10^−3^, respectively.

We performed variance analysis to quantify the extent to which the composite expression profiles are corrupted by the sample composition. In the composite data, ∼40 to 90% (*p*_rc_ < 2.7⋅10^−4^) of the variation is explained solely by the composition, as shown Fig. 3b. Interestingly, the effect varies between the genes. For instance, *KIF1A* expression has only 16% compositional effect, whereas *C1R* expression is explained by 77% by the composition, and the immune specific genes, e.g. *PPBP*, are more susceptible of having a cancer or fibroblast component. This suggests that the immune cell gene expression patterns are more dependent on the microenvironment composition than that of the other cell types. Consequently, previous analyses performed on patient tissue samples without accounting for the compositional factors likely remain useful, but may be biased towards findings in less compositionally affected gene sets.

When analyzing the PRISM-derived cell type specific profiles, only ∼0 to 15% of variation is explained by the composition, as shown in Fig. 3b. This indicates that PRISM can eliminate the confounding effect of composition variation in the decomposed signals and enrich the sample specific signal of the constituent cell types, as intended. Further, the remaining variation is captured by the cell type specific decomposed expression profiles (*p*_rc_ < 1.8 × 10^−8^; see Fig. 3b), suggesting that the signal passing trough to the decomposed cell types is both a significant explanatory factor and that it well captures the sample specificity of the original composite bulk sample.

We also verified that the cell type specificity of expression patterns is not limited to individual genes, but is reflected in pathway activity estimates as well. We derived gene set enrichment analysis (GSEA) scores ^20^ for the NCI Pathway Interaction Database (NCI-PID) ^21^ pathways from both the composite and decomposed data as shown in Fig. S8. While most of the differential pathway scores apper to originate from cancer cells, a significant effect is contributed by fibroblasts and immune cells depending on the pathway. For example, the NOTCH, FOXM1, and HIF1A pathway scores appear to originate from the cancer cells (<5.3% from other sources); RB1, PI3KCI, and FANCONI mostly from a combination of cancer and immune cells (<3.3%); FGF from cancer and stroma (<1.1%); and IL4 and IL23 mostly from immune cells (<2.9%), as shown in Fig. 3c and Fig. S9. Accordingly, the pathway scores using the decomposed profiles yield higher GSEA scores, indicating that the decomposition allows performing pathway analysis at a finer level of detail, by accounting for the compositional variation and removing the nuisance cell components, as suggested by Fig. 3b. The results were confirmed in the TCGA ovarian cancer dataset ^7^ (see Supplement).

Validation of the composition estimates. To verify that the composition is accurately estimated, we compared the PRISM estimates with estimates derived from whole-genome sequencing (WGS) data. Fig. S6a shows the correlation with ASCAT ^22^ purity estimates from the corresponding WGS data. The correlation is 77% (p-value *p*_lc_ < 6.3 ⋅ 10^−17^ for no linear correlation in a t-test). Further, we verified that the composition can be accurately estimated in other datasets and cancer types. Thus we applied PRISM on the TCGA ovarian cancer ^7^ bulk RNA sequencing data using our single-cell data; and to the TCGA skin cutaneous melanoma ^8^ bulk RNA sequencing data using the single-cell data from Tirosh et al. ^23^ and compared with the estimates from TCGA clinical data (immunohistochemistry) ^7, 8^, ABSOLUTE ^24^ (whole-genome sequencing), and LUMP ^6^ (methylation 450k array) from Aran et al. ^6^ (see Supplement). Finally, we verified that comparable composition estimates are obtained by using a single-cell panel derived from a different sequencing platform and when holding out the matching patients (see Supplement).

### Validation of cell type specificity of expression profiles

We performed RNA-ish experiments to verify that the PRISM decomposed profiles are indeed expressed differentially in cancer, stromal and immune cells. For this, we selected three genes for each cell type: *TRIM29*, *PARD6B*, *KIF1A* (cancer cell specific), *C1R*, *COL1A2*, *NAALADL2* (fibroblast), *RNASE6*, *GPR34*, and *C3AR1* (immune). The genes were selected to have high expression in the specific cell type (Fig. 3) and a significant difference between the complete response and progressive disease groups. As show in Fig. 4, RNA-ish validation was done with samples from seven HGSOC patients with matching bulk RNA-seq data, all nine genes being highly expressed in the PRISM predicted cell type (*p*_m_ < 10^−8^), except for *NAALADL2*.

**Figure 4.**
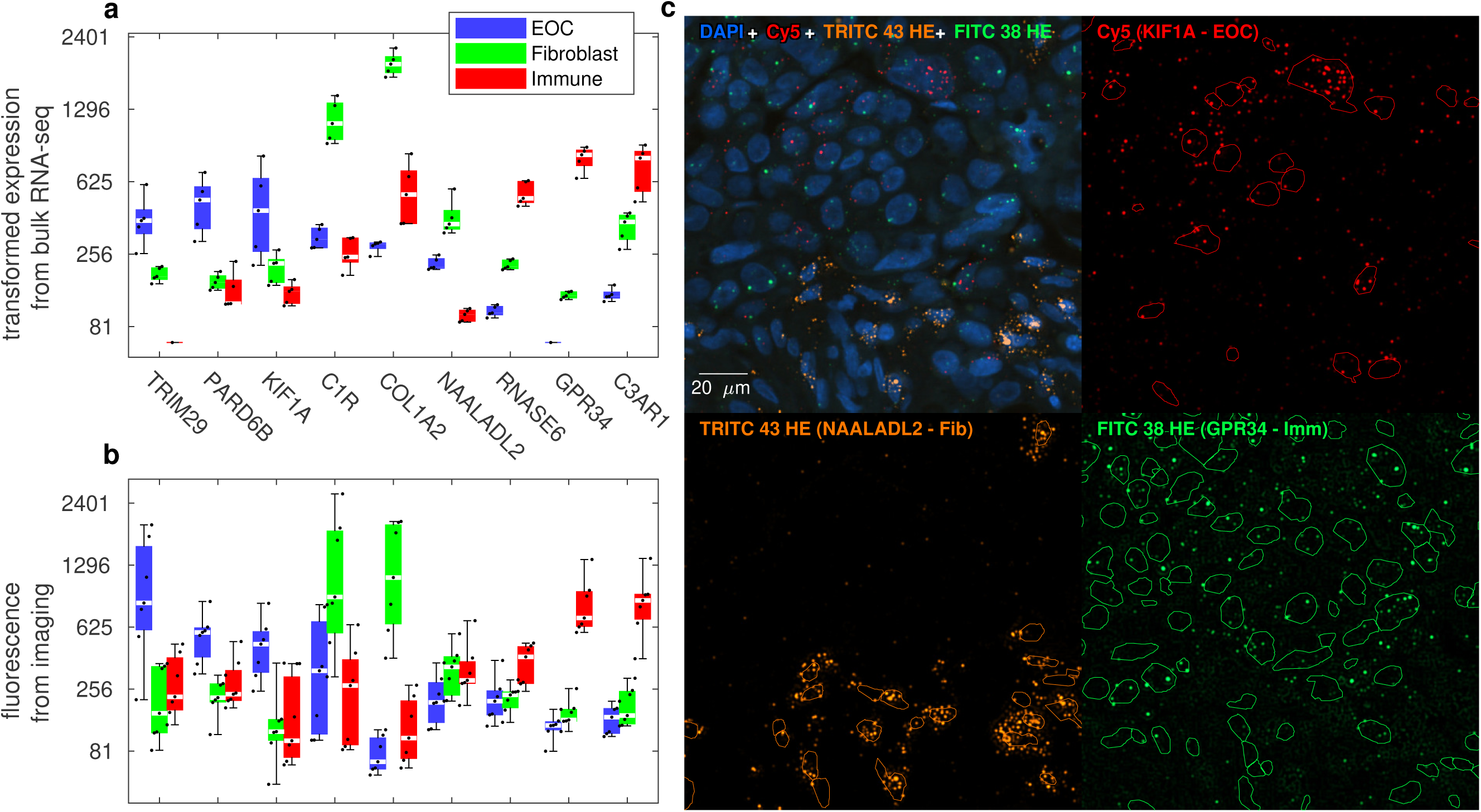
Validation of cell type specific expression patterns. a) Predicted expression level (scaled to match the RNA-ish experiment) from the decomposed bulk RNA samples grouped by the cell type for the seven matching samples seven patients and the selected nine genes. The box denotes first to third quartile, white bar median, and whiskers all data. Dots represent the samples, jittered by their rank. b) The corresponding quantified fluorescence from RNA-ish measurements. c) A region from the RNA-ish imaging, with split channels and our segmentation, showing the cell type specificity of the fluorescence tagged RNAs.

### Decomposed RNA profiles predict patient response

The PRISM analysis revealed several genes with expression level differences between complete response and the progressive disease patients groups, of which the most prominent are shown in Fig. S7. Cancer cell specific genes *TRIM29*, *PARD6B*, and *KIF1A* were found to be upregulated in the progressive disease group, while the fibroblast specific *C1R*, *COL1A2*, and *NAALADL2*, and immune specific *RNASE6*, *GPR34*, and *C3AR1* are downregulated in the progressive group (*p*_m_ < 6.5 ⋅ 10^−8^). In the RNA-ish data, the difference was significant for six genes (*KIF1A*, *C1R*, *COL1A2*, *RNASE6*, *GPR34*, and *C3AR1*), for *TRIM29* and *PARD6B* the trend was opposite, and for *NAALADL2* was inconclusive.

For *KIF1A*, *C1R*, and *GPR34*, we divided the 214 bulk RNA samples into the bottom 50% and top 50% groups by the expression level to predict the time to progression of the disease. As suggested by the differences between the complete response and progressive disease groups, we found that a high level of *KIF1A* in the cancer cell specific profile and low levels of *C1R* and *GPR34* in the fibroblast and immune specific profiles, respectively, confer less effective treatment and more rapid recurrence of the cancer. As shown in Fig. 5, this difference is not visible in the composite bulk signal. We verified that a similar association exists in the decomposed TCGA ovarian cancer ^7^ data for *KIF1A*, *C1R* (p-value *p*_h_ < 6.6 ⋅ 10^−5^ for equal hazards in a log-rank test), and *GPR34* (*p*_h_ = 0.07) regarding overall patient survival (see Fig. S15). While the trend is also visible in the composite bulk data for *KIF1A* (*p*_h_ = 9.2 ⋅ 10^−5^), the results for *C1R* are not (*p*_h_ = 0.20). In general, the survival associations are more significant for the decomposed data for the selected genes and at the whole-transcriptome scale in both ovarian cancer and in skin cutaneous melanoma (see Supplement).

**Figure 5.**
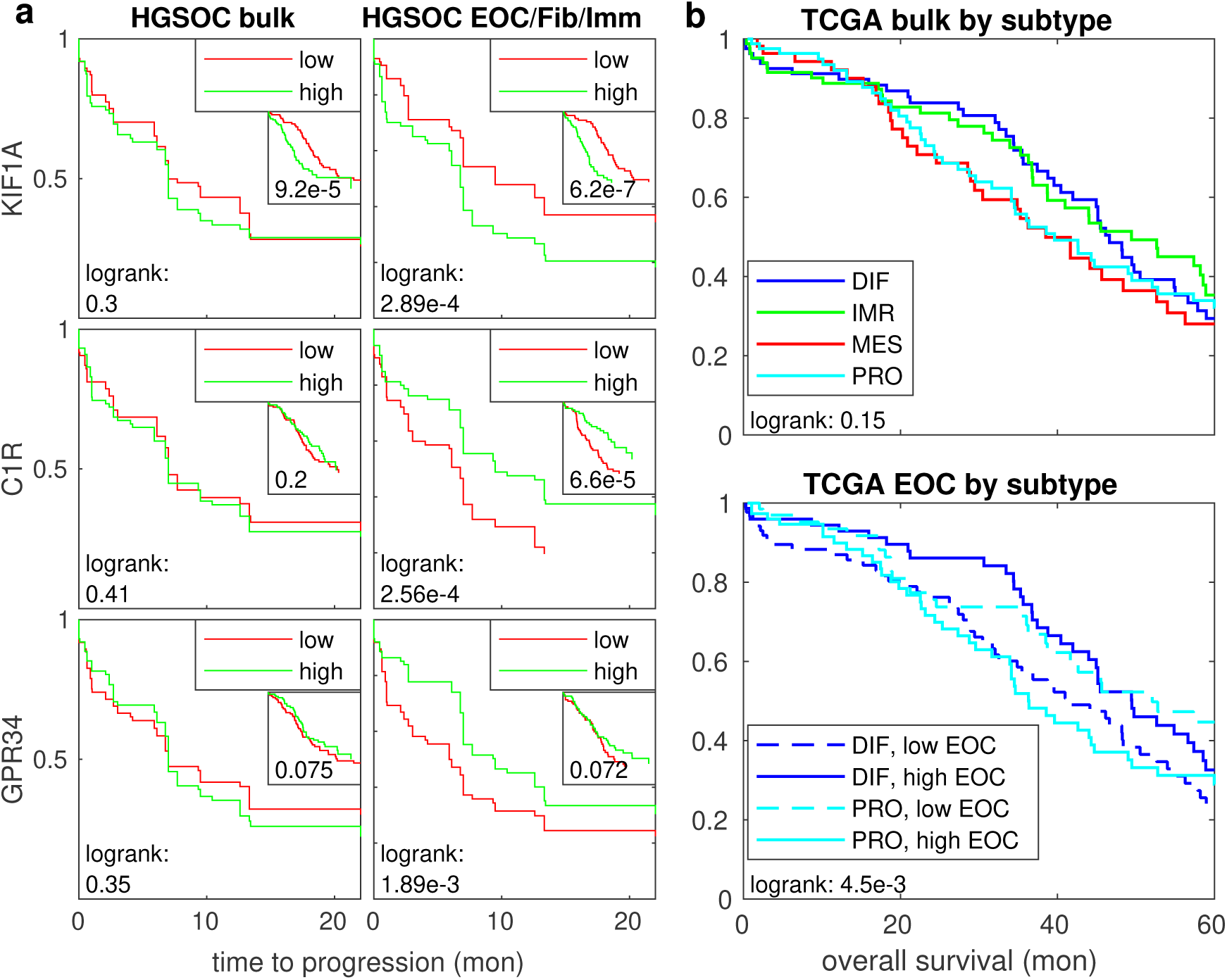
Survival association of the composite and decomposed RNA-seq data. a) Time to cancer progression between groups of samples with bottom 50% and top 50% expression of *KIF1A* (EOC), *C1R* (fibroblast), or *GPR34* (immune) when using the composite (left column) or the PRISM-derived cell type specific expression levels for the corresponding cell type (right column). The corresponding overall survival in the TCGA ovarian cancer ^7^ dataset are shown in the insets. b) Overall survival in the TCGA ovarian cancer dataset when grouped by the subtypes derived from composite bulk data (upper panel) or the PRISM-derived cancer cell specific signal (lower panel).

*KIF1A*, *C1R* and *GPR34* have not been associated with HGSOC survival earlier. *KIF1A* overexpression has been associated with cancer tissue in endometrial cancer ^25^ and it confers docetaxel resistance in breast cancer cell lines ^26^. Peptidase S1 protein family genes, such as *C1R*, are often expressed in the stroma and endothelium of various malignant tumors ^27, 28^, and are associated with innate immune response activation, inducing phagocytosis, among various functions ^28, 29^. *GPR34* is expressed primarily in specific immune cells ^30^ and has been shown to be required for adequate immune response in mice ^31^; it has been shown to be differentially expressed to the non-cancerous tissue in at least six different cancer types ^30^. The expression differences of these genes and their relevant function in other cancers warrants further study of these genes as prognostic and/or therapeutic targets.

Several studies have reported gene expression signatures in HGSOC and other cancers. As these are predominantly derived from bulk RNA-seq data, we tested their robustness in PRISM decomposed profiles. We derived HGSOC subtype estimates using the CLOVAR method ^32^, which classifies the samples into differentiated (DIF), immunoreactive (IMR), mesenchymal (MES), or proliferative (PRO) subtypes from both the composite and decomposed RNA profiles. Our results indicate that within the HGSOC subtypes, the IMR subtype highly depends on the immune cell frequency alone (77% correlation; see Fig. S10) and the MES subtype on fibroblasts (84%). DIF and PRO subtypes appear to originate from the cancer cells and feature weaker correlation with the composition (*p*_rc_ = 0.49 and 0.80, respectively), suggesting that these subtypes likely reflect phenotypic differences in the cancer cells, unlike the IMM and MES subtypes. The results were consistent between the our longitudinal and the TCGA ovarian cancer datasets.

In the TCGA dataset we found that deriving the subtypes in the absence of fibroblast and immune signals yields a significantly better separation in the overall survival (*p*_h_ < 4.5 ⋅ 10^−3^) than from the composite bulk data (*p*_h_ = 0.15), as shown in Fig. 5. To exclude the possibility that the gene expression signatures are unstable in HGSOC only, we analyzed gene expression signatures in TCGA skin cutaneous melanoma ^8^ dataset using the expression-derived subtypes ^8^. Here, the “immune” subclass reflects mostly immune cell frequency (77% correlation, *p*_rc_ < 3.0⋅10^−2^), while the MITF-low and keratin subtypes represent likely phenotypic differences between the cancer cells. Again, after removing the confounding immune component and the compositional variation, the patient classification predicts overall survival much better (*p*_h_ = 4.7 ⋅ 10^−3^ versus 0.022).

In general, our results indicate that some of the previously reported cancer subtypes obtained by clustering composite expression data are explained by the sample composition variation alone. This is in line with a previous report in head and neck cancer ^33^. While the composition may be indicative of patient survival (e.g. high immune content tends to correlate with better survival), our results show that the patient response and survival can be more accurately predicted by subtyping the cell type specific signals separately.

## Discussion

We developed a statistical framework, PRISM, for the analysis of heterogeneous RNA mixtures, and showed how it can be exploited for extracting the composition and the bulk-adapted whole-transcriptome profiles for each constituent cell type from each individual bulk RNA sample. By analyzing 214 longitudinal HGSOC samples, we showed that the tumor composition varies systematically with the treatment phase and the anatomical location, posing a challenge in personalized transcriptomic analysis. We showed that these challenges can be overcome with PRISM, which accurately estimates cell type specific expression profiles, which can serve as better predictors of patient response than bulk RNA-seq data. Importantly, analysis of 308 TCGA ovarian cancer, and 474 TCGA skin cutaneous melanoma samples agreed with these findings.

The main limitation of PRISM is that a heterogeneous sample of single-cell data from each cell type involved is required for consistent performance, which can be a problem if the reference and the analysis datasets are stratified according to different criteria. However, as we have shown, good performance can be expected without matching data as long as the single-cell data is not inherently biased. Further, as a statistical method, the expression profile estimates for infrequent cell types can be inaccurate, which may limit the analysis of (but not controlling for) stromal and immune cell subtypes. These points may warrant further investigation, but we expect that the issues are mitigated in the future as single-cell cataloging efforts move forward.

Precision oncology approach calls for methods that can exploit general statistical patterns in a cohort of a heterogeneous disease, but operate reliably at the individual patient level regardless of the evolving disease state, and adapt to the specifics of that patient, to which PRISM is a response regarding whole-transcriptome analysis of bulk samples. We believe PRISM has direct applications in analyzing transcriptomic data from other diseases that stem from heterogeneous causes and sampling setting, such as other cancer types, and that analysis methods for other genomic domains can benefit from the insights of our approach.

## Methods

### Patient and sample characteristics

The patient cohort consists of patients treated for ovarian or primary peritoneal HGSOC at Turku University Hospital between September 2010 to October 2018. All patients participating in the study gave written informed consent. The study and the use of all clinical material have been approved by The Ethics Committee of the Hospital District of Southwest Finland (ETMK) under decision number EMTK: 145/1801/2015.

We acquired 214 bulk RNA sequencing samples from 61 of the patients. Of these, 120 are primary (before chemotherapy), 60 interval (after chemotherapy), and 20 relapsed tumors (after being diagnosed as recurring). The samples are from primary ovarian tumors and various sites of intra-abdominal solid metastases and ascites fluid, as detailed in the analysis. Fig. 1a shows an overview of the sampling. Patient response is classified as complete response, partial response, stable disease, or progressive disease according to the RECIST criteria (version 1.1) ^34^. The sample collection and analysis is part of the HERCULES project (http://www.project-hercules.eu/).

### Single-cell RNA-seq sample preparation

Immediately after surgery, the HGSOC tumor specimens from our cohort were incubated overnight in a mixture of collagenase and hyaluronidase (Department of Pathology, University of Turku) to obtain single cell suspensions. Specimens were processed with a modified Fluidigm C1 protocol ^35^ or the standard Chromium Single Cell 3^′^ Reagent Kit v. 2.0 (10x Genomics) protocol for single-cell RNA sequencing with Illumina (HiSeq2000 for Fluidigm C1, HiSeq4000 or NextSeq for Chromium specimens (Jussi Taipale Lab, Karolinska Institute or Functional Genomics Unit, University of Helsinki).

We acquired 6,312 single cell profiles from 8 samples (from 7 patients and from various tissues) using the Chromium platform, and 347 cells from 8 samples (8 patients) using the Fluidigm single-cell sequencing platform. The latter were used for comparison purposes only. The single-cell samples were all matched to the bulk RNA samples but most bulk RNA samples remain unmatched.

### RNA-seq preprocessing

Bulk RNA sequencing reads were preprocessed using SePIA ^36^ pipeline within the Anduril framework ^37^. Read pairs were trimmed using Trimmomatic (version 0.33) ^38^ as follows: i) the first 12 bases were cropped due to uneven per base sequence content; ii) any leading bases with a quality score lower than 20 and any trailing bases with a quality score lower than 30 were removed; iii) the reads were scanned with a 5-base wide sliding window, cutting when the average quality per base drops below 20; iv) resulting sequences shorter than 20 bp were discarded. Trimmed reads were aligned to the GRCh38.d1.vd1 reference genome with GENCODE v25 annotation using STAR (version 2.5.2b) ^39^, allowing up to 10 mismatches, and all alignments for a read were output. Gene level effective counts (we found these to be more accurate than the raw read counts) were quantified using eXpress (version 1.5.1-linux x86 64) ^40^.

For the single-cell sequencing data, the raw base call (BCL) files were processed, including demultiplexation, alignment, barcode assignment and UMI quantification, with CellRanger (version 2.1.1) pipelines. The reference index was built upon the GRCh38.d1.vd1 reference genome with GENCODE v25 annotation. Single-cell transcriptomes were clustered using a shared nearest neighbor (SNN) modularity optimization based clustering algorithm implemented in Seurat (version 2.3.4) ^41^. PCA was selected as dimensional reduction technique in construction of SNN graph. Cell types were annotated based on acknowledged markers: epithelial cell markers: *WFDC2*, *PAX8*, *MUC16*, *EPCAM*, *KRT18*; stromal cell markers: *COL1A2*, *FGFR1*, *DCN*; immune cell markers: *CD14*, *CD79A*, *FCER1G*, *PTPRC*, *NKG7*, *CD3D*, *CD8A*.

### Modeling RNA expression data

We assume that latent cell type specific RNA counts *Z*_*ilj*_ exist, and can be approximated by a scaled Poisson distribution, i.e. 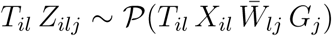, where the index 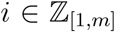 runs over the *m* genes, 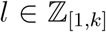 over the *k* cell types, and 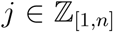 over the *n* samples, and *X*_*il*_ ∈ ℝ_≥0_ represents the cell type specific average expression profile, *T*_*il*_^−1^ ∈ ℝ_≥0_ is the dispersion (specific to each cell type and gene), 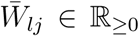 the convex composition, and *G*_*j*_ ∈ ℝ_≥0_ the sample specific scale factor. This approach allows capturing both biological and technical noise and accommodates either overdispersion (as is commonly observed in the data) and underdispersion (which improves stability under systematic errors) with respect to Poisson noise. The posterior of the observed 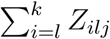 does not feature a closed form, but we show how to fit such models using an iterative algorithm (see Supplement). Unlike previous models ^42, 43^, we are not inconvenienced by the posterior tractability and account for the discrete and heteroscedastic nature of the data (i.e. genes and cell types are not equally reliable and informative), and freely varying dispersion confers estimator robustness.

### Decomposing bulk data using single-cell data

The model can be exploited for decomposing bulk data by considering a joint model on the bulk 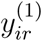 and single-cell data 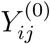. For each bulk, we assume that a cell type (and bulk specific) expression profiles (*X*_*il*_, *T*_*il*_) exist, as specified in the previous section, composing the bulk, i.e. 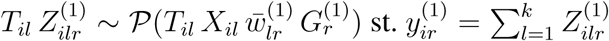, and being similar to the single-cell data, i.e. 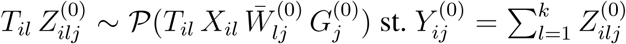, where *y*^(0)^ and *Y*^(1)^ and the single-cell and bulk data, respectively, *Z*^(0)^ and *Z*^(1)^ their latent random state, 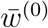 and 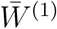 their convex composition, and *G*^(0)^ and *G*^(1)^ the sample scale factors. Again, *i* runs over the genes, *l* over the cell types, *j* over the single-cell profiles, and *r* over the bulk replicates (typically *r* = 1). As *T*_*il*_ can vary, the decomposition will weigh in the genes that are informative in discriminating the cell types. The cell type specific profiles in the bulk can be estimated as 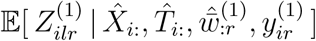 and the composition as 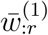, where 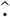 are the maximum likelihood estimates from the model fit. This process exploits all genes and all the single-cell data, but automatically weights down the non-relevant information across the two datasets, allowing the model to adopt to heterogeneous settings.

### Estimating scale factors

In mixtures, the scale factors are naturally estimated as part of the deconvolution process. Meanwhile, in pure (single-component) samples, the scale factors can be estimated by considering a fraction *α* of unperturbed genes, and finding an unperturbed common subprofile (*x*^(⊤)^, *t*^(⊤)^), i.e. 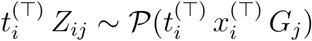 for some sparse set of genes 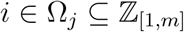 st. |Ω_*i*_| = *α m*, revealing a global relative scaling factor *G*_*j*_ for each single-cell sample.

### Discovering constituent phenotypes

In the decomposition, the composition *W*^(0)^ of the reference profiles (i.e. single-cell data) can be either preset or let vary freely. To facilitate more complex analyses, we also devised a hierarchical clustering process that exploits our model and reveals the cell types such that a single common bulk-independent consensus can be reached. The process is governed by the same model, i.e. 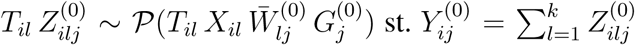, for the single-cell data *Y*^(0)^ but results in a binary composition 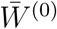 built up agglomeratively. This procedure is more stable against the multiple optima than an iterative algorithm, and allows selecting the optimal number of components using statistical means, such as Bayesian information criterion (BIC) unlike general-purpose clustering algorithms.

### RNA *in situ* hybridization

Formalin-fixed paraffin embedded (FFPE) tissue sections were analyzed using the RNAscope Multiplex Fluorescent Reagent Kit version 2 (#323100, Advanced Cell Diagnostics). We used catalog probes for the target RNAs for quantification and a positive and negative controls to verify good signal. The protocol is detailed in Supplement.

For fluorescence quantification, we used CellProfiler (version 3.1.8) ^44^ for segmentation, a Laplacian of Gaussian filter applied on a non-orthogonal basis projection for spot quantification, and cell classification based on fluorescence cosine-distance clustering (see Supplement).

## Acknowledgements

This work is supported in part by the European Union’s Horizon 2020 research and innovation programme under Grant Agreement No. 667403 for HERCULES (Comprehensive Characterization and Effective Combinatorial Targeting of High-Grade Serous Ovarian Cancer via Single-Cell Analysis), the Academy of Finland (Projects No. 292402, 325956, and 314395), the Sigrid Jusélius Foundation, and the Finnish Cancer Association. A.H. is funded by Academy of Finland grant No. 322927. We thank CSC—IT Center for Science Ltd. for compute resources. The results published here are in part based upon data generated by TCGA managed by the NCI and NHGRI. Information about TCGA can be found at https://cancergenome.nih.gov/. RNA-ish images were generated using 3DHISTECH Pannoramic 250 FLASH II digital slide scanner at Genome Biology Unit supported by HiLIFE and the Faculty of Medicine, University of Helsinki, and Biocenter Finland.

## Competing Interests

The authors declare that they have no competing financial interests.

## References

1. Schwartzberg, L., Kim, E. S., Liu, D. & Schrag, D. Precision oncology: Who, how, what, when, and when not? ASCO Educational Book 160–169 (2017).

2. Hanahan, D. & Weinberg, R. A. Hallmarks of cancer: the next generation. Cell 144, 646–674 (2011).

3. Aparicio, S. & Caldas, C. The implications of clonal genome evolution for cancer medicine. New Engl. J. Med. 368, 842–851 (2013).

4. Karczewski, K. J. & Snyder, M. P. Integrative omics for health and disease. Nat. Rev. Genetics 19, 299 (2018).

5. Lin, V. T. G. & Yang, E. S. The pros and cons of incorporating transcriptomics in the age of precision oncology. J. Natl. Cancer Inst. 111, 1–7 (2019).

6. Aran, D., Sirota, M. & Butte, A. J. Systematic pan-cancer analysis of tumour purity. Nat. Commun. 6, 8971 (2015).

7. The Cancer Genome Atlas Research Network. Integrated genomic analyses of ovarian carcinoma. Nature 474, 609–615 (2011).

8. The Cancer Genome Atlas Research Network. Genomic classification of cutaneous melanoma. Cell 161, 1681–1696 (2015).

9. Yoshihara, K. et al. Inferring tumour purity and stromal and immune cell admixture from expression data. Nat. Commun. 4, 2612 (2013).

10. Schelker, M. et al. Estimation of immune cell content in tumour tissue using single-cell RNA-seq data. Nat. Commun. 8, 1–12 (2018).

11. Sun, X., Sun, S. & Yang, S. An efficient and flexible method for deconvoluting bulk RNA-seq data with single-cell RNA-seq data. Cells 8, 1161 (2019).

12. Newman, A. M. et al. Robust enumeration of cell subsets from tissue expression profiles. Nat. Methods 12, 453–457 (2015).

13. Wang, X., Park, J., Susztak, K., Zhang, N. R. & Li, M. Bulk tissue cell type deconvolution with multi-subject single-cell expression reference. Nat. Commun. 10, 380 (2019).

14. Newman, A. M. et al. Determining cell type abundance and expression from bulk tissues with digital cytometry. Nat. Biotechnol. 37, 773–782 (2019).

15. Frishberg, A. et al. Cell composition analysis of bulk genomics using single-cell data. Nat. Methods 16, 327–332 (2019).

16. Torre, L. A. et al. Ovarian cancer statistics, 2018. CA Cancer J. Clin. 18, 284–296 (2018).

17. Ciriello, G. et al. Emerging landscape of oncogenic signatures across human cancers. Nat. Genet. 45, 1127–1133 (2013).

18. Ledermann, J. A. et al. Newly diagnosed and relapsed epithelial ovarian carcinoma: ESMO clinical practice guidelines for diagnosis, treatment and follow-up. Ann. Oncol. 24, vi24–vi32 (2013).

19. Hsiao, L.-L. et al. A compendium of gene expression in normal human tissues. Physiol. Genomics 7, 97–104 (2001).

20. Subramanian, A. et al. Gene set enrichment analysis: a knowledge-based approach for interpreting genome-wide expression profiles. Proc. Natl. Acad. Sci. U.S.A. 102, 15545–15550 (2005).

21. Schaefer, C. F. et al. PID: the Pathway Interaction Database. Nucl. Acids Res. 37, D674–D679 (2009).

22. Van Loo, P. et al. Allele-specific copy number analysis of tumors. Proc. Natl. Acad. Sci. U.S.A. 107, 16910–16915 (2010).

23. Tirosh, I. et al. Dissecting the multicellular ecosystem of metastatic melanoma by single-cell rna-seq. Science 352, 189–196 (2016).

24. Carter, S. L. et al. Absolute quantification of somatic DNA alterations in human cancer. Nat. Biotechnol. 30, 413–421 (2012).

25. Wong, Y. F. et al. Identification of molecular markers and signaling pathway in endometrial cancer in Hong Kong chinese women by genome-wide gene expression profiling. Oncogene 26, 1971–1982 (2007).

26. De, S., Cipriano, R., Jackson, M. W. & Stark, G. R. Overexpression of kinesins mediates docetaxel resistance in breast cancer cells. Cancer Res. 69, 8035–8042 (2009).

27. Bulla, R. et al. C1q acts in the tumour microenvironment as a cancer-promoting factor independently of complement activation. Nat. Commun. 7, 10346 (2015).

28. Reis, E. S., Mastellos, D. C., Ricklin, D., Mantovani, A. & Lambris, J. D. Complement in cancer: untangling an intricate relationship. Nat. Rev. Immunol. 18, 5–18 (2018).

29. Markiewski, M. M. & Lambris, J. D. Is complement good or bad for cancer patients? a new perspective on an old dilemma. Trends Immunol. 30, 286–292 (2008).

30. Schoneberg, T., Meister, A., Jaroslawna Bernd Knierim & Schulz, A. The G protein-coupled receptor GPR34 — the past 20 years of a grownup. Pharmacol. Therapeut. 189, 71–88 (2018).

31. Liebscher, I. et al. Altered immune response in mice deficient for the G protein-coupled receptor GPR34. J. Biol. Chem. 286, 2101–2110 (2011).

32. Verhaak, R. G. et al. Prognostically relevant gene signatures of high-grade serous ovarian carcinoma. J. Clin. Invest. 123, 517–525 (2013).

33. Puram, S. V. et al. Single-cell transcriptomic analysis of primary and metastatic tumor ecosystems in head and neck cancer. Cell 171, 1611–1624 (2017).

34. Eisenhauer, E. A. et al. New response evaluation criteria in solid tumours: Revised RECIST guideline (version 1.1). Eur. J. Cancer 45, 228–247 (2009).

35. Islam, S. et al. Quantitative single-cell RNA-seq with unique molecular identifiers. Nat. Methods 11, 163–166 (2014).

36. Icay, K. et al. SePIA: RNA and small RNA sequence processing, integration, and analysis. BioData Min. 9, 20 (2016).

37. Cervera, A. et al. Anduril 2: upgraded large-scale data integration framework. Bioinformatics 35, 3815–3817 (2019).

38. Bolger, A. M., Lohse, M. & Usadel, B. Trimmomatic: a flexible trimmer for Illumina sequence data. Bioinformatics 30, 2114–2120 (2014).

39. Dobin, A. et al. STAR: ultrafast universal RNA-seq aligner. Bioinformatics 29, 15–21 (2013).

40. Roberts, A. & Pachter, L. Streaming fragment assignment for real-time analysis of sequencing experiments. Nat. Methods 10, 71–73 (2013).

41. Satija, R., Farrell, J. A., Gennert, D., Schier, A. F. & Regev, A. Spatial reconstruction of single-cell gene expression data. Nat. Biotech. 33, 495 (2015).

42. Robinson, M. D., McCarthy, D. J. & Smyth, G. K. edger: a bioconductor package for differential expression analysis of digital gene expression data. Bioinformatics 26, 139–140 (2010).

43. McCarthy, D. J., Chen, Y. & Smyth, G. K. Differential expression analysis of multifactor RNA-Seq experiments with respect to biological variation. Nucl. Acids Res. 40, 4288–4297 (2012).

44. Kamentsky, L. et al. Improved structure, function and compatibility for CellProfiler: modular high-throughput image analysis software. Bioinformatics 27, 1179–1180 (2011).

