## Supplementary material for "PRISM: Recovering cell type specific expression profiles from composite RNA-seq data"

### Derivation of the models and methods

**Models and the underlying theory.** We assume that the distribution of each RNA (isoform) in the cells can be approximated by a scaled Poisson distribution. This is a valid model if the RNA are produced and degraded according to a Poisson process, or a similar process with smaller or greater dispersion [S1–S3]. We denote this by:

$$\mathcal{D}_{il}^{R} \doteq \tilde{\mathcal{P}}(\lambda_{il}, {t_{il}}^{-1}) \tag{S1}$$

where $\lambda_{il} \in \mathbb{R}_{\geq 0}$ represents the average expression of the $i$:th gene in the $l$:th cell type and $t_{il} \in \mathbb{R}_{\geq 0}$ is a precision parameter, $t_{il} = 1$ for Poisson process, and $t_{il} > 1$ ($t_{il} < 1$) for more (less) deterministic. Generally, heterogeneous convex mixtures of Poisson random variables are well modeled by $t_{il} < 1$. Multimodal distributions cannot be represented, but this is not necessary, as multimodality can be achieved by using multiple components in what follows. Any (positive) distribution can be matched up to the two first moments (mean and variance), and performing model fitting automatically selects the best approximation (in K-L divergence sense). Note that $t_{il}$ models physiological noise, caused by phenotypic differences such as concentration of different metabolites, and is immune to sampling noise such as that arising from cDNA amplification and quantification (modeled later).

Physically, the shape parameter ${t_{il}}^{-1}$ can be thought to represent a burst size, i.e. quantity of items produced simultaneously uniformly in time at rate $\lambda_{il}$. However, in general, it is more useful to think this as how correlated the production events are: ${t_{il}}^{-1} > 1$ implies that multiple are produced at once (or the production is correlated), which is disordely if the interproduction intervals are random; while $t_{il} < 1$ corresponds to an assembly line like process, single events built up in substeps piecemeal, which is orderly.

Next, we assume the $j$:th sample contains $N_{lj}$ cells (generally not known and depends on factors like volume of initial sample content to be sequenced and amplification factor) of the $l$:th cell type. Following this, the (unobserved) number of RNA isoforms of type $i$ specific to this cell type in the $j$:th sample is:

$$\begin{aligned} \tilde{Z}_{ilj} &= \sum_{w=1}^{N_{lj}} R_{ilj}^{(w)} = \tilde{\mathcal{P}}(\lambda_{il}\, N_{lj}, {t_{il}}^{-1}) \\ R_{ilj}^{(w)} &\sim \mathcal{D}_{il}^{R} \quad \text{iid (over } w) \end{aligned} \tag{S2}$$

If the population is sampled using a random process, such as RNA sequencing [S1], with efficiency $\eta_j \in [0, 1]$, the (latent) isoform counts at quantification for each cell type are:

$$\begin{aligned} Z_{ilj} \mid \tilde{Z}_{ilj} &\sim \mathcal{B}(\tilde{Z}_{ilj}, \eta_j) \\ Z_{ilj} &\approx \tilde{\mathcal{P}}(\underbrace{\lambda_{il}}_{\doteq X_{il}}\ \underbrace{N_{lj}\,\eta_j}_{\doteq W_{lj}},\ \underbrace{(1-\eta_j) + \eta_j\, {t_{il}}^{-1}}_{\approx {T_{il}}^{-1}}) \end{aligned} \tag{S3}$$

where the approximation is exact for Poisson $\tilde{Z}_{ilj}$, exact to the first two moments at minimum (i.e.

unbiased with respect to mean and variance), and exact asymptotically as $\lambda_{il}\, N_{lj}\, \eta_j \to \infty$.

The sampled counts $Z_{ilj}$ are Poisson iff $t_{il} = 1$, in which case the conversion efficiency plays no role. Otherwise, the precision of the latent counts $T_{ilj}$ is controlled by both $t_{il}$ and $\eta_j$, and only depends on $i, l$ if the conversion efficiencies $\eta_j$ are approximately constant. Note that the exact value of the precision (as quantified later) is rather meaningless, as it depends on $\eta_j$, but does not otherwise depend on $j$. As expected, the conversion efficiency interpolates the gene specific noise–for low efficiency ($\eta_j \to 0$) the observations are Poissonian, as the random sampling noise dominates, and for high efficiency ($\eta_j \to 1$) this approaches the biological variation of the gene ${t_{il}}^{-1}$, as the sampling is perfect. Note that, regardless of the regime the amount of RNA $N_{lj}$ plays no role on the noise, so $T_{ilj}$ can be mostly considered independent of the sample. Regardless, some regularizing assumption is needed for the model to be identifiable, and we find this the most useful, as variation over either genes or cell types are expected to dominate.

The scaled Poisson approach allows modeling both under- and overdispersion, which can result from correlations, (random) RNA dropout, and the model representing additional mixtures, such as cell subtypes and cellular heterogeneity. Our dispreference for the more commonly used negative binomial distribution [S2] stems from two facts: i) the scaled Poisson is more tractable analytically, while still allowing accurate modeling of overdispersion; ii) as underdispersion is allowed, the model is more stable numerically under systematic errors (e.g. from read mapping and isoform quantification). We note that modeling RNA sequencing data using a negative binomial is generally not physically motivated, but only found to fit the measurements [S1].

A measurement in a bulk sample quantifies the sum of the sampled counts, parametrized by $\mathbf{X}$, $\mathbf{T}$, and $\mathbf{W}$:

$$Y_{ij} = \sum_{l=1}^{k} Z_{ilj} \mathrel{\dot\sim} \mathcal{M}\Big( ((X_{il})_{i=1}^{m})_{l=1}^{k}, ((T_{il})_{i=1}^{m})_{l=1}^{k}, ((W_{lj})_{l=1}^{k})_{j=1}^{n} \Big) \tag{S4}$$

where $\mathbf{X}$ represents the cell type specific expression profiles, $\mathbf{T}$ the cell type specific precision (inverse noise) profiles, and $\mathbf{W}$ the composition, as defined above. The posterior (i.e. the distribution of $Y_{ij}$ does not have a closed form representation, and cannot be approximated as it might be multimodal, but this is not problematic (as we show later). With a single active component, the model generalizes to single-cell data.

**Parameter estimation.** We use the latent variables $Z_{ilj}$ and the observations $Y_{ij}$ from above:

$$\begin{aligned} Z_{ilj} &\sim \tilde{\mathcal{P}}(X_{il} W_{lj}, T_{il}) \\ Y_{ij} &= \sum_{l=1}^{k} Z_{ilj} \end{aligned} \tag{S5}$$

which allows the parameters $\mathbf{X}$, $\mathbf{T}$, and $\mathbf{W}$ to be conveniently estimated in maximum likelihood sense (i.e. we choose the parameters that most likely generate the data) using an iterative expectation maximization (EM) algorithm. The algorithm consist of an E-step, where the sufficient statistics of $Z_{ilj}$ are acquired, given the current set of parameters and the data; followed by the M-step, which in turn updates the parameter set by maximizing the expected likelihood over $Z_{ilj}$ [S4]. It can be shown that an increase in the expected likelihood implies and increase in the likelihood of the full model, implying that a local maximum can be found by iterating the steps under some mild regularity conditions [S4].

For the M-step we seek to improve the expected latent likelihood given a current set of parameters $\theta^{(0)} = (\mathbf{X}^{(0)}, \mathbf{T}^{(0)}, \mathbf{W}^{(0)})$:

$$\begin{aligned}
&\mathbb{E}_{\mathbf{Z}\,|\,\theta^{(0)},\mathbf{Y}}\left[\log g(\,\mathbf{Z}\,|\,\mathbf{X},\mathbf{T},\mathbf{W}\,)\,|\,\theta^{(0)},\mathbf{Y}\,\right] \\
= \quad &\sum_{i=1}^{m}\sum_{l=1}^{k}\sum_{j=1}^{n}\Bigg(\log T_{il} \;+\; T_{il}\;\mathbb{E}[\,Z_{ilj}\,|\,\theta^{(0)},\mathbf{Y}\,]\;\log(T_{il}\,X_{il}\,W_{lj}) \;+ \\
&\qquad\qquad - \mathbb{E}[\,\log\Gamma(T_{il}\,Z_{ilj}+1)\,|\,\theta^{(0)},\mathbf{Y}\,] - T_{il}\,X_{il}\,W_{lj}\Bigg)
\end{aligned} \tag{S6}$$

which is maximized at:

$$\begin{aligned}
X_{il} &= \frac{\sum_{j'=1}^{n}\mathbb{E}[\,Z_{ilj'}\,|\,\theta^{(0)},\mathbf{Y}\,]}{\sum_{j'=1}^{n}W_{lj'}} \\
{T_{il}}^{-1} &\approx \frac{1}{n}\sum_{j'=1}^{n}\frac{\overbrace{\mathbb{E}[\,Z_{ilj'}\,|\,\theta^{(0)},\mathbf{Y}\,]\Big(\,\mathbb{E}[\,Z_{ilj'}\,|\,\theta^{(0)},\mathbf{Y}\,]-X_{il}\,W_{lj'}\Big)}^{\text{unexplained variability over the } n \text{ samples}} + \overbrace{\mathbb{C}\mathrm{ov}[\,Z_{ilj'}\,|\,\theta^{(0)},\mathbf{Y}\,]}^{\text{mixing variability}}}{X_{il}\,W_{lj'}} \\
W_{lj} &= \frac{\sum_{i'=1}^{m}T_{i'l}\;\mathbb{E}[\,Z_{i'lj}\,|\,\theta^{(0)},\mathbf{Y}\,]}{\sum_{i'=1}^{m}T_{i'l}\,X_{i'l}}
\end{aligned} \tag{S7}$$

where the approximation uses the truncation of:

$$\begin{aligned}
\log\Gamma(z+1) &= z\,\log(z) - z + \frac{1}{2}\,\log(z) + \frac{1}{2}\,\log(2\,\pi) + \mathcal{O}(1/z) \\
\mathbb{E}[\,Z\,\log(Z)\,] &= \mathbb{E}[Z]\,\log(\mathbb{E}[Z]) + \frac{1}{2}\,\frac{\mathbb{C}\mathrm{ov}[Z]}{\mathbb{E}[Z]} + \mathcal{O}((Z-\mathbb{E}[Z])^3)
\end{aligned} \tag{S8}$$

which is exact for Poisson ($T_{il} = 1$) and exact asymptotically for high expression levels $X_{il}\,W_{il} \to \infty$ or for low noise $T_{il} \to \infty$. We note that data that are both low mean and high noise possess low Fisher information (i.e. are close to missing data), so they must have a small impact on a maximum-likelihood procedure anyway, rendering the approximation very useful. (The first one is what is well known as the Stirling's approximation, and the second is a second-order Taylor expansion around the mean of a random variable.)

To obtain the expectations relevant to Eq. (S7), consider:

$$
\begin{aligned}
Z_{ilj} &\sim \mathcal{P}(X_{il}\, W_{lj}) \\
Z_{ilj} \,|\, \sum_{l'=1}^{k} Z_{ilj} = Y_{ij} &\sim \mathcal{B}(\, Y_{ij}, \frac{X_{il}\, W_{lj}}{\sum_{l'=1}^{k} X_{il'}\, W_{l'j}}\,)
\end{aligned} \tag{S9}
$$

and $K \,|\, y \sim \mathcal{B}(y, p)$ implies $\mathbb{E}[\, K \,|\, y\,] = p\, y$ and $\mathbb{C}\mathrm{ov}[\, K \,|\, y\,] = p\,(1-p)\, y$. Or more generally, when $K_l \sim \tilde{\mathcal{P}}(\lambda_l, t_l)$, $\sum_{l=1}^{k} K_l = y$, and $t_l \neq 1$:

$$
\begin{aligned}
\mathbb{E}[\, K_l \,|\, y\,] &\approx \frac{\lambda_l}{\lambda_\Sigma}\, y \\
\mathbb{C}\mathrm{ov}[\, K_l \,|\, y\,] &\approx \frac{\lambda_l}{\lambda_\Sigma} \left(1 - \frac{\lambda_l}{\lambda_\Sigma}\right) \left(\frac{1}{\lambda_\Sigma} \sum_{l'=1}^{k} \frac{\lambda_{l'}}{t_{l'}}\right) y \\
\text{where} \quad \lambda_\Sigma &\doteq \sum_{l'=1}^{k} \lambda_{l'}
\end{aligned} \tag{S10}
$$

which is exact for all $t_l$ equal (including but not limited to Poisson and $t_l \to \infty$) and otherwise asymptotic at $y \to \infty$. This is a generalization of a multinomial distribution where the items are contageous (i.e. ${t_l}^{-1}$-clusters of items are drawn iid with probability of $\frac{\lambda_l}{\lambda_\Sigma}$ to a total of $y$ items), which again highlights the physical role of $t_l$. Again, the posterior lacks closed form, but the above moments can be derived from the recurrence:

$$
\mathbb{P}[\, K_1 = y_1, \cdots, K_k = y_k\,] = \sum_{l=1}^{k} \frac{\lambda_l}{\lambda_\Sigma}\; \mathbb{P}[\, K_1 = y_k, \cdots, K_l = y_l - {t_l}^{-1}, \cdots, K_k = y_k\,] \tag{S11}
$$

on a $k$-dimensional semi-open integer lattice $(y_1, \cdots, y_k) \geq \mathbf{0}$, which is a generalization of the multinomial generator obtained at $t_l = 1$. The small $y$ behavior is intricate due to discretization effects, and hence an asymptotic approximation is used. Again, small counts feature low Fisher information (e.g. zero for $y = 0$), so the inaccuracies tend to occur in uninformative regions.

Finally, we can outline the parameter estimation algorithm:

1. Choose initial $\mathbf{X}$, $\mathbf{T}$, and $\mathbf{W}$. The choice is not important if all classes are observed (e.g. decomposing a bulk with single cell data from all cell types), as e.g. in some cases the objective can be shown to be convex, but the initial parameter must not be on the boundary (i.e. $X_{il} = 0$, $T_{il} = 0$, or $W_{lj} = 0$). We begin with the profiles $\mathbf{X} \leftarrow \mathbf{1}$ uniform, the shape $\mathbf{T} \leftarrow \mathbf{1}$ Poisson, and the composition $\mathbf{W} \leftarrow \mathbf{1}$ to uniform.

2. Find the latent mean and covariance of $\mathbf{Z} \mid \mathbf{X}, \mathbf{T}, \mathbf{W}, \mathbf{Y}$ using Eq. (S10)

3. Estimate new $\mathbf{X}$, $\mathbf{T}$ using Eq. (S7).

4. Update the latent variables $\mathbf{Z} \mid \mathbf{X}, \mathbf{T}, \mathbf{W}$ using Eq. (S10) and update $\mathbf{W}$ using Eq. (S7). The equations in Eq. (S7) are mutually recursive on $((\mathbf{X}, \mathbf{T}), \mathbf{W})$, so two EM stages are required.

5. Repeat until convergence as indicated by the gradient norm of Eq. (S6)

**Decomposition of bulk RNA sequencing data.** The above models can be used to decompose bulk RNA samples using single-cell data as follows. The algorithm simultaneously finds the cell-type specific expression patterns composing each sample, and the sample composition (i.e. fraction of each cell type), as illustrated in Figure S1. Pre-labeled single-cell data are used to guide the composition. The labels can be obtained through clustering or classification using marker genes (we have used the latter approach), or could be derived from validated cell lines. (Technically the method also works without single-cell data, performing blind decomposition, or with unknown single-cell labeling inferred in the process, but we did not evaluate the performance.)

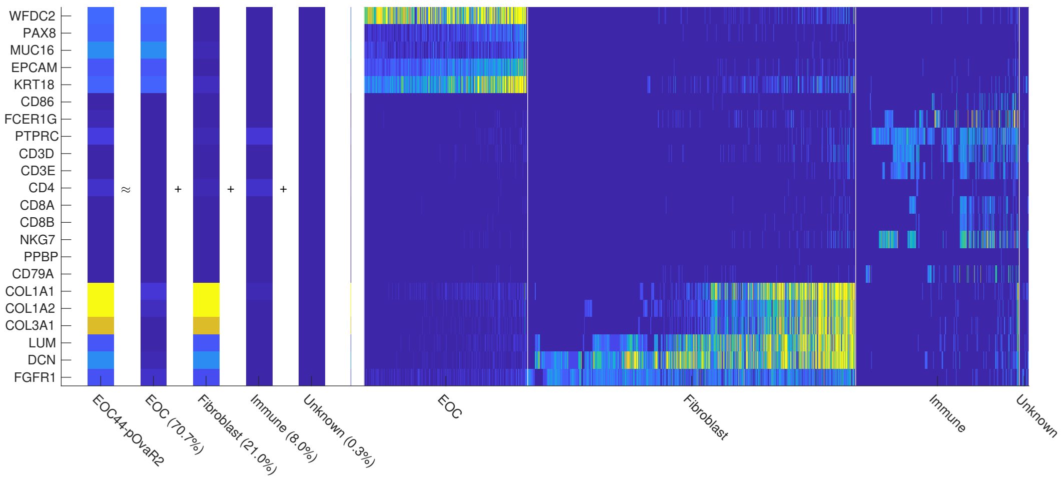


**Figure S1** Decomposing bulk RNA sequencing data into cell specific components. Left bars: each bulk sample is composed of cell-type specific expression patterns (unique to each sample), mixed in unknown proportions. Our decomposition reveals both the expression patterns and the composition with the aid of labeled single-cell data. Right bars: single-cell expression data where cell-type specific expression patterns are apparent.

The decomposition is based on the following model:

$$\begin{aligned} \mathbf{Y}^{(0)} &\sim \mathcal{M}(\mathbf{X}, \mathbf{T}, \mathbf{W}^{(0)}) \\ \mathbf{y}^{(1)} &\sim \mathcal{M}(\mathbf{X}, \mathbf{T}, \mathbf{w}^{(1)}) \end{aligned} \tag{S12}$$

where $\mathbf{Y}^{(0)} \in \mathbb{R}_{\geq 0}^{m \times n}$ are the single-cell expression patterns (reference), the $m$ rows representing genes and the $n$ columns the individual cells, $\mathbf{W}^{(0)} \in \mathbb{R}_{\geq 0}^{k \times n}$ are the known weights for the single cells, the $k$ rows representing the decomposed cell types, $\mathbf{y}^{(1)} \in \mathbb{R}_{\geq 0}^{m \times r}$ are the bulk expression patterns, the $r$ columns representing the replicates, $\mathbf{W}^{(1)} \in \mathbb{R}_{\geq 0}^{k \times r}$ are the (unknown) weights for the bulk sample, representing the composition, and $\mathbf{X} \in \mathbb{R}_{\geq 0}^{m \times k}$ are the (unknown) cell-type specific expression profiles and $\mathbf{T} \in \mathbb{R}_{\geq 0}^{m \times k}$ its precision matrix, capturing both physiological and technical noise, as discussed earlier. The weight matrices also contain the sample gains, the scaling or batch factors, (i.e. $\mathbf{W}^{(i)} = \bar{\mathbf{W}}^{(i)}\mathbf{G}^{(i)}$) where $\bar{\mathbf{W}}^{(i)} \in \mathbb{R}_{\geq 0}^{k\times n}$ are the (column) normalized weights summing to unity and $\mathbf{G}^{(i)} \in \mathbb{R}_{\geq 0}^{n\times n}$ is a diagonal matrix of the sample gains). The bulk sample gains $\mathbf{G}^{(1)}$ are naturally estimated but $\mathbf{G}^{(0)}$ are not. A solution for finding $\mathbf{G}^{(0)}$ is developed in the next section. (Note that often $r = 1$ and the process is repeated for each bulk sample, which results in a separate (bulk specific) expression patterns $\mathbf{Z}$, but this need not be the case.). An illustration how the data and variables are related is shown in Figure S2, and a plate diagram for the model is shown in Figure S3.

The unknown parameters, $\mathbf{X}$, $\mathbf{T}$, and $\mathbf{w}^{(1)}$, are solved as specified in the previous section. Finally, the cell type specific expression patterns in the bulk sample are estimated as:

$$\mathbb{E}[\,\mathbf{Z}\,|\,\hat{\mathbf{X}}, \hat{\mathbf{T}}, \mathbf{W}^{(0)}, \hat{\mathbf{w}}^{(1)}, \mathbf{Y}^{(0)}, \mathbf{y}^{(1)}\,] \tag{S13}$$

the composition as $\hat{\bar{\mathbf{w}}}^{(1)}$, and the sample gains as $\hat{\mathbf{G}}^{(1)}$, where $\hat{\cdot}$ are the maximum likelihood parameter estimates resulting from the EM algorithm.

The described decomposition procedure is readily available through the `prism-decom` driver in our implementation (`https://bitbucket.org/anthakki/prism/`).

**Estimating sample gains.** Resolving cell type specific gain factors is possible when the cell types are present in a mixture, where extrinsic scaling factors such as amount of initial RNA material or the amplification factor are constant. However, for single-component data such as single-cells from various platforms, this cannot be done, as the intrinsic and extrinsic scaling factors factor

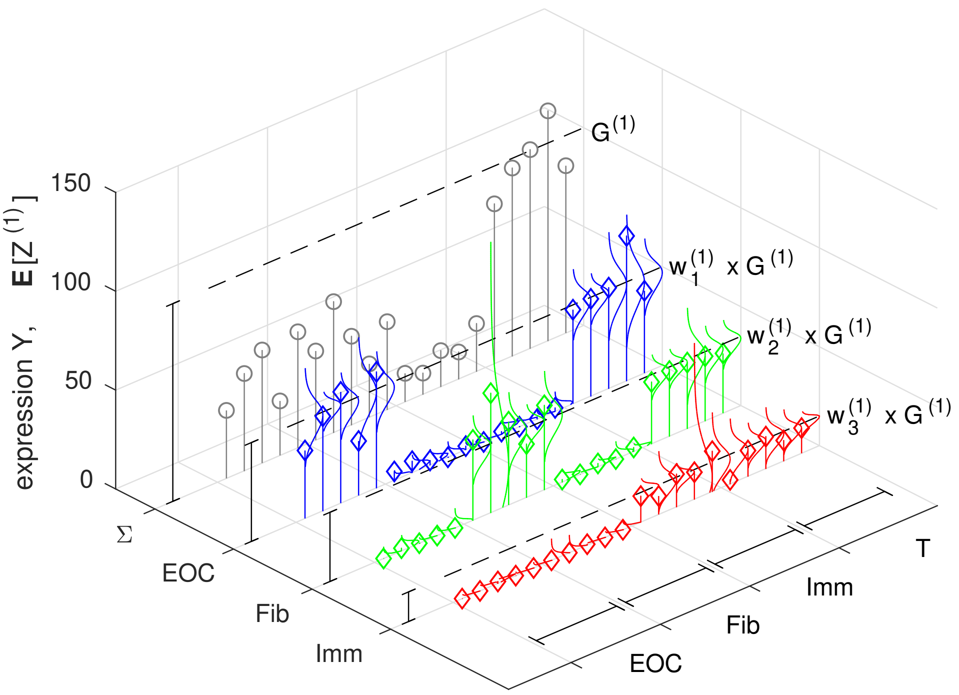


**Figure S2** Illustration how the bulk variables are related. Circles denote observed data, diamonds (inferred) estimates, and the dashed lines the (inferred) scaling. For each sample, there exist a readout profile ($\Sigma$) and cell-type specific profiles, whose expression level is defined by the scaling factor $G^{(1)}$ and the sample composition $w^{(1)}$. The scaling factors are revealed by the unchanged genes ($\top$) and the composition by the cell type specific genes (clustered by specificity, horizontal segments). Each gene features a cell type specific expression level and variability, and as shown, the distribution (vertical curves) can vary from geometric-like to a normal-like. Single-cell data is modeled with the same model, but with a single component, and the generating parameters $X_{il}, T_{lj}$ are shared.

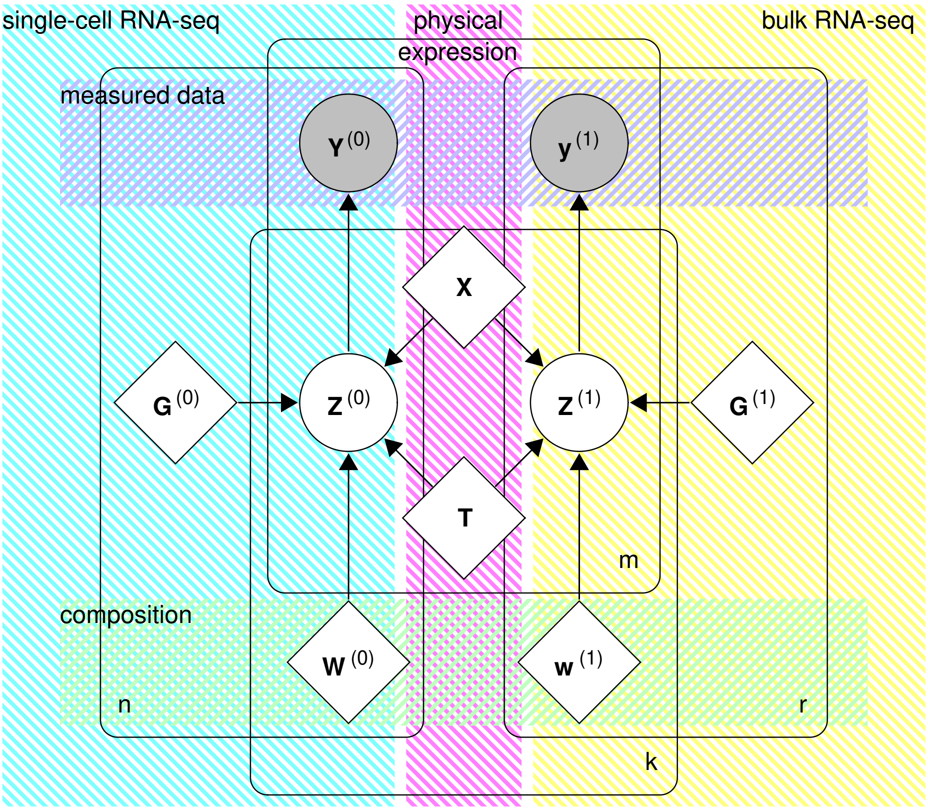


**Figure S3** Plate diagram for decomposing bulk data with $m$ genes, $k$ cell types, $j$ single-cell samples and $r$ bulk replicates. Circles represent random variables (gray observed and white latent) and diamonds (unknown) parameters. $X_{il}$ represent the expected (average) cell type specific expression profiles, ${T_{ij}}^{-1}$ the associated variability (or how it scales with the expression level), parametrizing the latent bulk $Z^{(1)}_{ilr}$ and single-cell $Z^{(0)}_{ilj}$ cell type specific expression read counts. The composition $\bar{W}^{(0)}_{lj}$, $\bar{w}^{(1)}_{lr}$ and scaling factors $G^{(0)}_j, G^{(1)}_r$ also affect the latent state $Z^{(0)}_{ilj}$, $Z^{(1)}_{ilr}$ which generate the mixture RNA readouts $Y^{(0)}_{ij}, y^{(1)}_{ir}$.

into a single scale element. This is a recognized problem, and can be accounted for in downstream analysis, such as differential expression testing [S2], but it poses a challenge for the decomposition, as unequal scaling factors can bias the compositional interpretation.

A naive total count (sum of reads) normalization is likely to be problematic in this context, as e.g. the cancer cells proliferating much faster are likely to feature much higher global expression level. Fortunately, alternative strategies like using control genes for the normalization exist [S5].

Next, we show how our framework can be utilized to resolve overcome this challenge. By assuming that there exist some (unknown) set of genes, a proportion of $\alpha$ in each sample, whose expression is not (significantly) affected by the cell type. These genes could be the control genes from prior knowledge, but need not to be so and need not to be known. In fact, the genes need not to be even common in all samples (cell types).

Consider the following model for the single-cell data $\mathbf{Y}^{(0)}$:

$$\mathbf{Y}^{(0)} \sim \mathcal{M}(\,\mathbf{x}^{(\top)}, \mathbf{t}^{(\top)}, \mathbf{G}^{(0)}) \quad \text{with weights} \quad \mathbf{U}^{(0)} \tag{S14}$$

where $\mathbf{x}^{(\top)} \in \mathbb{R}_{\geq 0}^{m\times 1}$ and $\mathbf{t}^{(\top)} \in \mathbb{R}_{\geq 0}^{m\times 1}$ represent the unknown globally unchanged expression profile, $\mathbf{G}^{(0)} \in \mathbb{R}_{\geq 0}^{1\times n}$ the unknown single-cell specific gains, and $\mathbf{U}^{(0)} \in \{0,1\}^{m\times n}$ is an unknown binary indicator for the unchanged genes in each sample with a density $\alpha$.

The following outlines an EM procedure for estimating the global unchanged expression profiles and the single-cell specific gains:

1. Acquire an initial solution by solving $\mathbf{x}^{(\top)}$ and $\mathbf{t}^{(\top)}$ for $\mathbf{G}^{(0)} = \mathbf{1}$ and $\mathbf{U}^{(0)} = \mathbf{1}$, and then solve for $\mathbf{G}^{(0)}$, using Eq. (S7). Note that $Z_{ilj} \,|\, \mathbf{x}^{(\top)}, \mathbf{t}^{(\top)}, \mathbf{G}^{(0)}, \mathbf{Y} = Y_{ij}$, as there is only a single component.

2. Compute the membership probabilities using Eq. (S6), select $\alpha\, m$ least-likely outlier genes in each sample, and set weights $\mathbf{U}^{(0)}$ accordingly. This can be done in $\mathcal{O}(m\, n)$ time.

3. Estimate new parameters using an EM iteration using Eq. (S7) by leaving out the perturbed terms as indicated by $\mathbf{U}^{(0)}$.

4. Repeat until converge.

This process is similar in rationale than the trimmed mean of M-values (TMM) method [S6] in that the scaling factors are estimated from partial input data, but also accounts for the heteroscedasticity. Our method also works around small $\mathbf{Y}$, unlike TMM, which is chaotic around $Y_{ij} = 0$. The trimming will mitigate this, but globally low expression levels can cause a problem, which is more susceptible in occurring in the single-cell data than in bulks. Our method also accommodates multiple samples without specifying a pairwise ordering (which is suboptimal and depends how the samples are scheduled) and without having to compute all the $\mathcal{O}(n^2)$ pairs and having an explicit merging procedure.

Finally, we note that the estimates $\mathbf{x}^{(\top)}$ and $\mathbf{t}^{(\top)}$ might not be as useful by themselves as one could intially think. When $\mathbf{U}^{(0)}$ is sparse, then $\mathbf{x}^{(\top)}$ and $\mathbf{t}^{(\top)}$ might have many unmeaningful elements, which do not contribute to the model fit. Secondly, the trimming will underestimate variance, so $\mathbf{t}^{(\top)}$ are overestimated, with respect to the true underlying unchanged profile (but not with respect to the trimmed set of samples). Similarly, $\mathbf{x}^{(\top)}$ can feature a bias when the distribution is skewed and gets trimmed.

This and some of the other normalization methods are available using the `prism-gain` driver in our implementation (`https://bitbucket.org/anthakki/prism/`).

**Clustering single-cell data.** The putative cell types for the bulk components can be automatically discovered from the single-cell data. This could use any general purpose clustering algorithm, but we can exploit the proposed probabilistic model for the purposes, which is expected to be more accurate for sequencing data.

One way of exploiting our model would be fitting it for various values of $k$, potentially forcing sparsity on $\mathbf{W}$ in the EM loop, which would result in a k-means-like clustering algorithm. However, this can be computationally expensive, and a good result is hard to guarantee, due to the numerous optima (unlike in the decomposition, where the single-cell data weight out the alternative optima). For this reason, we adopt a full agglomerative hierarchical clustering procedure, which features consistent performance and runtime. Clustering using more specialized models for RNA-seq data have been proposed previously, but they tend to be of the k-means type [S7,S8].

To derive the clustering algorithm, we assume $k = 1$, as each single-cell is supposedly a pure signal composed out of a single (unknown) cell type, and that the gains $w_j$ are known (as determined in the previous step or by alternative means). A natural information-theoretic cost $c(p)$

for a cluster $p$ with the set of items $J_p$ can be defined:

$$\begin{aligned}
c(p) &\doteq \sum_{j\in J_p} \max_{\mathbf{x}_j,\mathbf{t}_j} \ell_{\mathcal{M}(\mathbf{x}_j,\mathbf{t}_j,w_j)}(\mathbf{y}_j) - \max_{\mathbf{x}_p,\mathbf{t}_p} \sum_{j\in J_p} \ell_{\mathcal{M}(\mathbf{x}_p,\mathbf{t}_p,w_j)}(\mathbf{y}_j) \\
&\approx \sum_{j\in J_p}\sum_{i=1}^{m} -\frac{1}{2}\log\frac{\hat{t}_j^{(p)}}{\hat{t}_j^{(j)}} - \hat{t}_j^{(p)}\, y_{ij}\,\log\frac{\hat{x}_i^{(p)}\, w_j}{y_{ij}} + \hat{t}_j^{(p)}\left(\hat{x}_i^{(p)}\, w_j - y_{ij}\right) \\
\text{where}\quad \hat{x}_i^{(p)} &\doteq \left(\sum_{j\in J_p} w_j\right)^{-1} \sum_{j\in J_p} y_{ij} \\
\hat{t}_i^{(p)\,-1} &\doteq \frac{1}{|J_p|}\left(\sum_{j\in J_p} \frac{y_{ij}\left(y_{ij} - \hat{x}_i^{(p)}\, w_j\right)}{\hat{x}_i^{(p)}\, w_j}\right)
\end{aligned} \tag{S15}$$

where $(\hat{x}_i)_{i=1}^m$ and $(\hat{t}_i)_{i=1}^m$ are the maximum likelihood (estimated) parameters for the cluster, $\hat{t}_j^{(j)}$ representing the precision of a singleton, which is a constant $\hat{t}_j^{(j)} \to \infty$ but the choice does not affect the clustering. Specifically, $c(p) = -\log\Lambda \geq 0$ where $\Lambda$ is the likelihood ratio between a one cluster model and a $|J_p|$ cluster model (i.e. each sample in a separate cluster) for the items in $J_p$, and is analogous to the sum of squared error to cluster centroid for a homoscedastic normal model (as in e.g. k-means or Ward's clustering [S9]). The approximation uses Stirling's approximation as discussed earlier, and the cost is exact for all $\hat{t}_j = 1$ (Poisson) or $\hat{t}_j \to \infty$ (low noise).

Analogous to Ward's method [S9], we define a distance that seeks for the least decrement (as opposed to the best fitting model, which results in an uninteresting tree) in the cluster likelihood:

$$\begin{aligned}
&d(p,q) \doteq c(p\cup q) - c(p) - c(q) \\
\approx &\underbrace{\sum_{i=1}^{m}\left(-\frac{|J_{p\cup q}|}{2}\log \hat{t}_i^{(p\cup q)} - \hat{t}_i^{(p\cup q)}\,\Sigma_{y_i}^{(p\cup q)}\log\frac{\Sigma_{y_i}^{(p\cup q)}}{\Sigma_{w}^{(p\cup q)}} + \hat{t}_i^{(p\cup q)}\,\Sigma_{z_i}^{(p\cup q)}\right)}_{\tilde{c}(p\cup q)} - \tilde{c}(p) - \tilde{c}(q)
\end{aligned}$$

$$\begin{aligned}
\text{where}\quad &\Sigma_{y_i}^{(p\cup q)} \doteq \sum_{j\in J_p\cup J_q} y_{ij} \quad\text{and}\quad \Sigma_{q_i}^{(p\cup q)} \doteq \sum_{j\in J_p\cup J_q} y_{ij}\,y_{ij}\,{w_j}^{-1} \qquad \text{(S16)}\\
&\Sigma_{w}^{(p\cup q)} \doteq \sum_{j\in J_p\cup J_q} w_j \quad\text{and}\quad \Sigma_{z_i}^{(p\cup q)} \doteq \sum_{j\in J_p\cup J_q} y_{ij}\,\log(\,y_{ij}\,{w_j}^{-1}\,)\\
&\hat{t}_i^{(p\cup q)\;-1} = \frac{1}{|J_{p\cup q}|}\left(\Sigma_{w}^{(p\cup q)}\,\Sigma_{q_i}^{(p\cup q)}\;\Sigma_{y_i}^{(p\cup q)\;-1} - \Sigma_{y_i}^{(p\cup q)}\right)
\end{aligned}$$

which can be updated in $\mathcal{O}(m)$ time, and is a reducible distance [S10]. This distance is not ultrametric, but the cumulative distance recovers the negative log-likelihood ratio at the maximum likelihood estimate $-\log\Lambda_k$, which is monotonic increasing, zero at the leaves if $\hat{t}_j^{(j)} = 1$ is chosen (representing a Poisson prior with infinitesimal weight), and can be visualized as a dendrogram, as shown in Figure S4.

A hierarchical clustering tree can be efficiently incrementally built using the reciprocal nearest neighbors (RNN) algorithm [S10]. The algorithm to build the hierarchical clustering tree in $\mathcal{O}(m\,n^2)$ time is as follows (merging and distance computation are $\mathcal{O}(m)$ time, as shown above):

1. Begin with each of the $n$ single-cell samples in its own cluster

2. Starting from cluster $p$, move to cluster $q$, its nearest neighbor (i.e. $\forall r : d(p,q) \leq d(p,r)$), unless $p$ is the nearest neighbor of $q$. Initial $p$ can be arbitrarily chosen, but after each merge the chain must be backtracked to last free $p$ for the search to be amortized $\mathcal{O}(n)$ [S10].

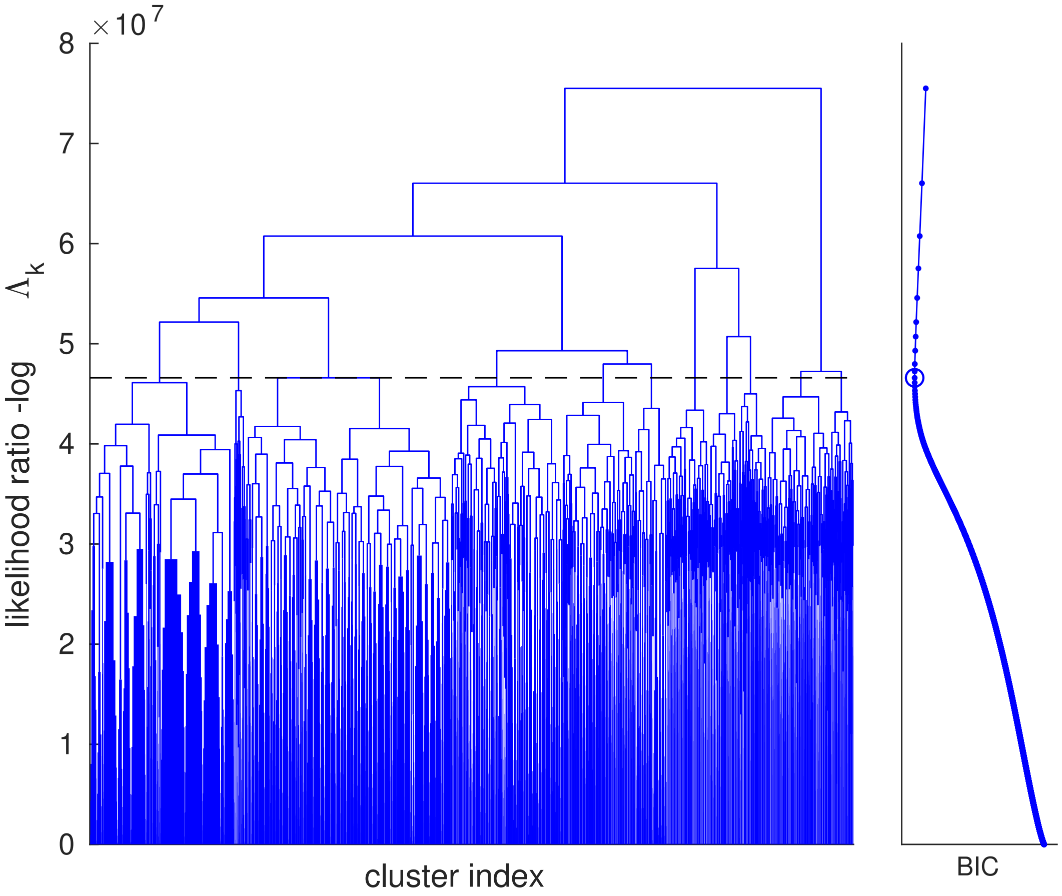


**Figure S4** Dendrogram of the hierarchical clustering tree obtained by clustering the $6{,}312$ single-cell samples with $58{,}037$ genes each in the Chromium ovarian cancer dataset. The plot is truncated at $500$ branches, the remaining subtrees appearing as solid rectangles. The curve on the right shows BIC for each node in the tree, with the minimum BIC value marked by a circle, suggesting $11$ clusters, and the dashed line indicates this cut.

3. Merge clusters $p$ and $q$ using Eq. (S16)

4. Terminate when only one cluster remains; otherwise, move to step 2

By the virtue of the probabilistic model specified above, the tree can be cut by statistical means, such as using Bayesian information criterion (BIC) or a likelihood ratio test (with a pre-specified significance level for false cluster formation). That is, the number of clusters $\hat{k}$ is selected such that:

$$\hat{k} \doteq \operatorname*{argmax}_k \left\{ \hat{\ell}_k - \frac{1}{2}\,\lambda\,k \right\} = \operatorname*{argmax}_k \left\{ \log \Lambda_k - \frac{1}{2}\,\lambda\,k \right\} \tag{S17}$$

where $\hat{\ell}_k$ is the log-likelihood for a $k$ cluster model, $\Delta = 2\,m$ the difference in the model degrees of freedom between $k$ cluster and $k-1$ cluster models, and $\lambda$ is the penalty for the model complexity. Specifically, $\lambda = \log(m\,n)\,\Delta$ for BIC, $\lambda = 2\,\Delta$ for Akaike information criterion (AIC), or $\lambda$ : $\mathbb{P}[\,\chi^2_\Delta \geq \lambda\,] = \alpha$ where $\chi^2_\Delta$ is a chi-squared variate with $\Delta$ degrees of freedom for likelihood ratio tests between successive models at significance level $\alpha$ (using an asymptotic approximation using Wilks' theorem). We prefer BIC, as it is consistent (i.e. asymptotically correct) but the others have different attractive properties [S11].

The single-cell clustering and model selection is available using the `prism-clust` command in our implementation (`https://bitbucket.org/anthakki/prism/`), and the results can be imported and visualized in R or MATLAB using a combination of the built-in hierarchical clustering tools.

### Supplementary methods

**Preprocessing for the single-cell Fluidigm RNA sequencing data.** For the Fluidigm single-cell RNA sequencing data, quality of the single-end reads were estimated with FastQC (`https:`

//www.bioinformatics.babraham.ac.uk/projects/fastqc/) and the reads were trimmed with Trimmomatic [S12] with the same steps as for bulk RNA sequencing reads. Trimmed reads were aligned to the GRCh38.d1.vd1 reference genome with GENCODE v25 annotation using STAR [S13], allowing up to 3 mismatches, and only uniquely mapped reads were output. UMI deduplication were performed using UMI-tools (version 0.2.3) [S14] with the “directional-adjacency” method. Gene level counts were quantified from deduplicated BAM files using featureCounts (version 1.5.1) [S15].

**Whole-genome sequencing analysis.** Raw WGS sequencing reads were subject to quality and adapter trimming using Trimmomatic (version 0.33) [S12] (minimum length 50, headcrop 5, sliding window $5:20$). Trimmed reads were then mapped to human reference genome GRCh38 using bwa mem (version 0.7.12) [S16] with default parameters. Mapped reads in BAM format were marked for duplicates using PICARD (version 2.6) tools (https://broadinstitute.github.io/picard/). Base quality recalibration was performed afterwards using the Genome Analysis toolkit (GATK; version 3.7; https://software.broadinstitute.org/gatk/). Copy number alterations were detected using ASCAT (version 2.3) [S17] following the AscatNGS workflow for sequencing data (https://github.com/cancerit/ascatNgs). For samples lacking a matched normal control, we used an unmatched normal control for computation of logR values, and we used the “ascat.predictGermlineGenotypes” function for B-allele frequency computation. ASCAT allows the joint estimation of ploidy and aberrant cell fraction (cellularity) assuming the ploidy of the non-aberrant cells is 2, revealing the fraction of cancer cells.

**RNA *in situ* hybridization and imaging.** RNA *in situ* hybridization was performed on fresh 4.5 µm formalin-fixed paraffin embedded (FFPE) tissue sections using RNAscope Multiplex Fluorescent Reagent Kit Version 2 (#323100, Advanced Cell Diagnostics) for target detection according to the manual. The experiments were performed in sets-of-three (RNASE6-TRIM29-C1R, PARD6B-COL1A2-C3AR1, and NAALADL2-GRP34-KIF1A) for each sample, where each RNA product was quantified using fluorescein (FITC 38 HE), Cyanine 3 (TRITC 48 HE), or Cyanine 5 (Cy5) tags, respectively, in a single experiment.

Firstly, tissue sections were baked for 1 h at 60 °C, then deparaffinized and treated with hydrogen peroxide for 10 min at room temperature (RT). Target retrieval was performed for 15 min at 98 °C, followed by protease plus treatment for 15 min at 40 °C. All probes (Table S1) were hybridized for 2 h at 40 °C followed by signal amplification and developing of HRP channels was done according to manual. TSA Plus fluorophores fluorescein (1:750 dilution), Cyanine 3 (1:1500 dilution) and Cyanine 5 (1:3000 dilution) (NEL744001KT, Perkin Elmer) were used for signal detection. The sections were counterstained with DAPI and mounted with ProLong Gold Antifade Mountant (P36930, Invitrogen). Tissue sections were scanned using 3DHISTECH Pannoramic 250 FLASH II digital slide scanner at Genome Biology Unit (Research Programs Unit, Faculty of Medicine, University of Helsinki, Biocenter Finland) using $1 \times 20$ magnification with extended focus and seven (7) focus levels.

**Quantitative analysis of RNA *in situ* hybridization.** We used the IdetifyPrimaryObjects component of CellProfiler (version 3.1.8) [S18] for segmentation in the DAPI staining. Non-default param-

**Table S1** List of probes used for the RNA *in situ* experiments. The table lists the probes used in the measurements and their RNAscope catalog number.

| Probe | Catalog number |
|---|---|
| Hs-TRIM29-C1 | 319871 |
| Hs-COL1A2-C1 | 432721 |
| Hs-GPR34-C1 | 521021 |
| Hs-C1R-C2 | 508951-C2 |
| Hs-C3AR1-C2 | 461101-C2 |
| Hs-KIF1A-C2 | 488931-C2 |
| Hs-NAALADL2-C3 | 540991-C3 |
| Hs-PARD6B-C3 | 426971-C3 |
| Hs-RNASE6-C3 | 476471-C3 |
| 3-plex negative control probe, dapB | 320871 |
| 3-plex positive control probe, POLR2A, PPIB, UBC | 320861 |

eters were as follows: typical diameter $18$ to $56$ pixels (determined experimentally), thresholding using adaptive Otsu's method (i.e. local not global), clumped object detection and splitting using shape (i.e. not intensity), and low-resolution speedups were disabled.

Cross-channel fluorescence bleed was reduced by finding a suitable basis near for the intensity data of all pixels near the principal axes using power iteration. This procedure resembles principal component analysis (PCA), but orthogonality is not enforced, so the expression signals can be correlated (as opposed to being uncorrelated by construction in PCA). Note that this is done at the pixel level, not on the cellular cellular level which is used for quantification, so at cellular level the channels can very well be correlated when multiple spots are localized inside one cell.

Next, the fluorescence intensity signal was quantified using the negative response of a Laplacian of Gaussian filter with standard deviation of unity (spot diameter of $\sim 2.36$ pixels at half the maximum). The value was tuned manually, and the kernel width roughly corresponds to the diameter of an observed RNA spot in our images. This procedure filters out background variations and cellular autofluorescence, leaving intensity blobs of the specified size. The remaining intensity was averaged in each segmented cell, the cells were weighted by area, and clustered into three $(3)$ classes using cosine distance. Each cluster is labeled using the nearest intensity axis.

In general, the expression cannot be assumed to be directly proportional to the quantified intensity, due to e.g. cellular autofluorescence, the properties of the different filters, and nonlinear response in the imaging procedure (gamma). This is handled in the post-processing, as we use distribution-free methods for statistical analysis.

### Supplementary results

**Major constituent cell types can be automatically discovered from the single-cell data.** In the previous section, we also showed how the developed models and methods can be used for automatically discovering the constituent cell types. We also experimented with this clustering to label the single-cell data. Figure S5 shows the clustering for the Chromium 6,312 Chromium single-cell samples with BIC criterion for model selection (resulting in 11 clusters).

The matching with our manually-assisted labeling of cell types was $97.7\%$, significantly better than would be expected by chance (p-value of $1.1 \times 10^{-4}$ in $m \times n$ Fisher's exact test for the cell type vs. cluster label crosstabulation below). The clustering further suggests that while the cancer cells of the patients tend to cluster separately, the fibroblasts and immune cells are less susceptible for this, suggesting that even limited panels of the neighboring cell types can improve the decomposition. Even if the clustering tree was cut to four clusters (as in our manual labeling), the matching is good ($73.6\%$; p-value $1.2 \times 10^{-4}$), the error being mostly due to the merged fibroblast-immune cell cluster of clusters 9 to 11 from below. Figure S6d shows a comparison of the estimated composition with our manually-assisted labeling versus the nearest BIC-derived labels, indicating that all cell types are accurately discovered (linear correlation of $98.3\%$, p-value $< 10^{-8}$ in a t-test for zero correlation).

**Validation of composition estimate accuracy.** We verified the accuracy of the PRISM composition estimates by various means. First, we compared the estimated tumor cell fraction to estimates derived using corresponding WGS data using ASCAT [S17], as shown in Figure S6a. ASCAT is

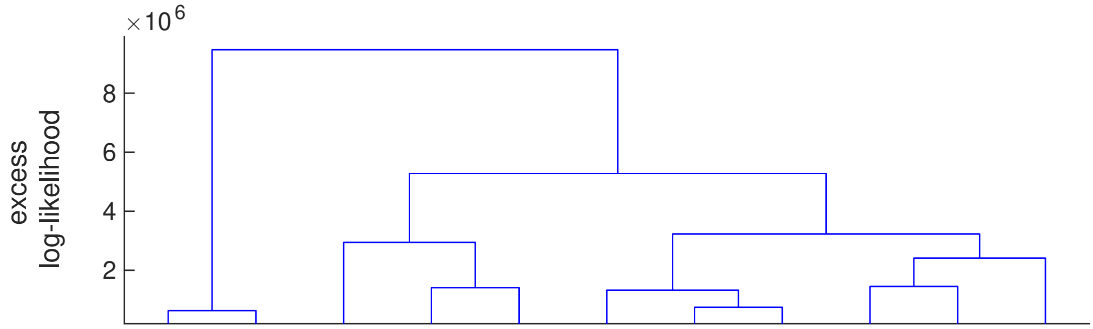


| | 1 | 2 | 3 | 4 | 5 | 6 | 7 | 8 | 9 | 10 | 11 |
|---|---|---|---|---|---|---|---|---|---|---|---|
| EOC | 521 | 195 | 347 | 246 | 227 | 14 | 2 | 0 | 0 | 0 | 1 |
| Fibroblast | 0 | 0 | 0 | 3 | 0 | 821 | 580 | 1 | 6 | 6 | 1697 |
| Immune | 0 | 0 | 0 | 1 | 3 | 5 | 13 | 258 | 1186 | 86 | 1 |
| Unknown | 0 | 0 | 0 | 0 | 1 | 86 | 3 | 0 | 0 | 0 | 2 |

| | 1 | 2 | 3 | 4 | 5 | 6 | 7 | 8 | 9 | 10 | 11 |
|---|---|---|---|---|---|---|---|---|---|---|---|
| EOC293_pAsc | 521 | 195 | 0 | 0 | 3 | 0 | 0 | 0 | 0 | 0 | 0 |
| EOC568_pTubL | 0 | 0 | 347 | 0 | 0 | 126 | 2 | 0 | 2 | 0 | 0 |
| EOC568_iOvaR | 0 | 0 | 0 | 0 | 0 | 5 | 0 | 6 | 3 | 0 | 942 |
| EOC649_iMes | 0 | 0 | 0 | 0 | 10 | 474 | 4 | 97 | 512 | 28 | 0 |
| EOC736_iOvaL | 0 | 0 | 0 | 0 | 2 | 10 | 0 | 19 | 36 | 34 | 745 |
| EOC781_pOme | 0 | 0 | 0 | 0 | 215 | 27 | 2 | 2 | 1 | 2 | 13 |
| EOC940_pPer | 0 | 0 | 0 | 249 | 0 | 283 | 590 | 130 | 7 | 15 | 0 |
| EOC1030_pAsc | 0 | 0 | 0 | 1 | 1 | 1 | 0 | 5 | 631 | 13 | 1 |

**Figure S5** Hierarchical clustering of $6{,}312$ single-cell profiles. The top panel shows the hierarchical clustering tree, cut to $11$ clusters using BIC, while the tables show how the different cell types (with manually assisted labeling) and the samples distribute to the different clusters.

susceptible for giving erroneously a purity of unity in some cases, while our method can estimate also the purity in lower purity settings. To counter this, we used the non-aberrant sample indicator of ASCAT to discard WGS samples whose tumor cell fraction cannot be reliably estimated. Second, we verified that the estimates remain consistent while holding out the matching single-cell data, as shown in Figure S6b. Third, we used a separate single-cell reference of $347$ single-cells using the Fluidigm platform (as opposed to the $6{,}312$ cells from the Chromium platform) to show that the estimates remain consistent across the single-cell sequencing platforms, as shown in Figure S6c. The composition estimates for TCGA ovarian cancer [S19] and skin cutaneous melanoma [S20] datasets were also compared with estimates derived from immunohistochemistry, genomic, and methylation data [S21], as shown in Figure S13 and Figure S16, respectively.

**Generalizability to non-matching patient data.** Next, we verified that the results generalize to the setting where matching single cell data is not available. Our dataset contained $4$ samples (from $4$ patients) featuring both single-cell and bulk data derived from the same tumor, which were used in a separate analysis with the matching single-cell data held out sequentially for each sample. As shown in the top right panel of Figure S6, we found the compositional estimates to remain consistent (linear correlation of $99.4\%$, p-value $< 5.0 \times 10^{-15}$), suggesting that the method operates robustly regardless if matched data is present or not (provided that sufficient breadth exists in the single cell reference such that population variability is not underestimated). We note that proportions of each cell type are robustly estimated, not just that of the cancer cells, which was already found accurate using the WGS comparison. We also found that, in agreement, the decomposed expression profiles remain consistent as well (rank correlation of $99.5\%$, p-value

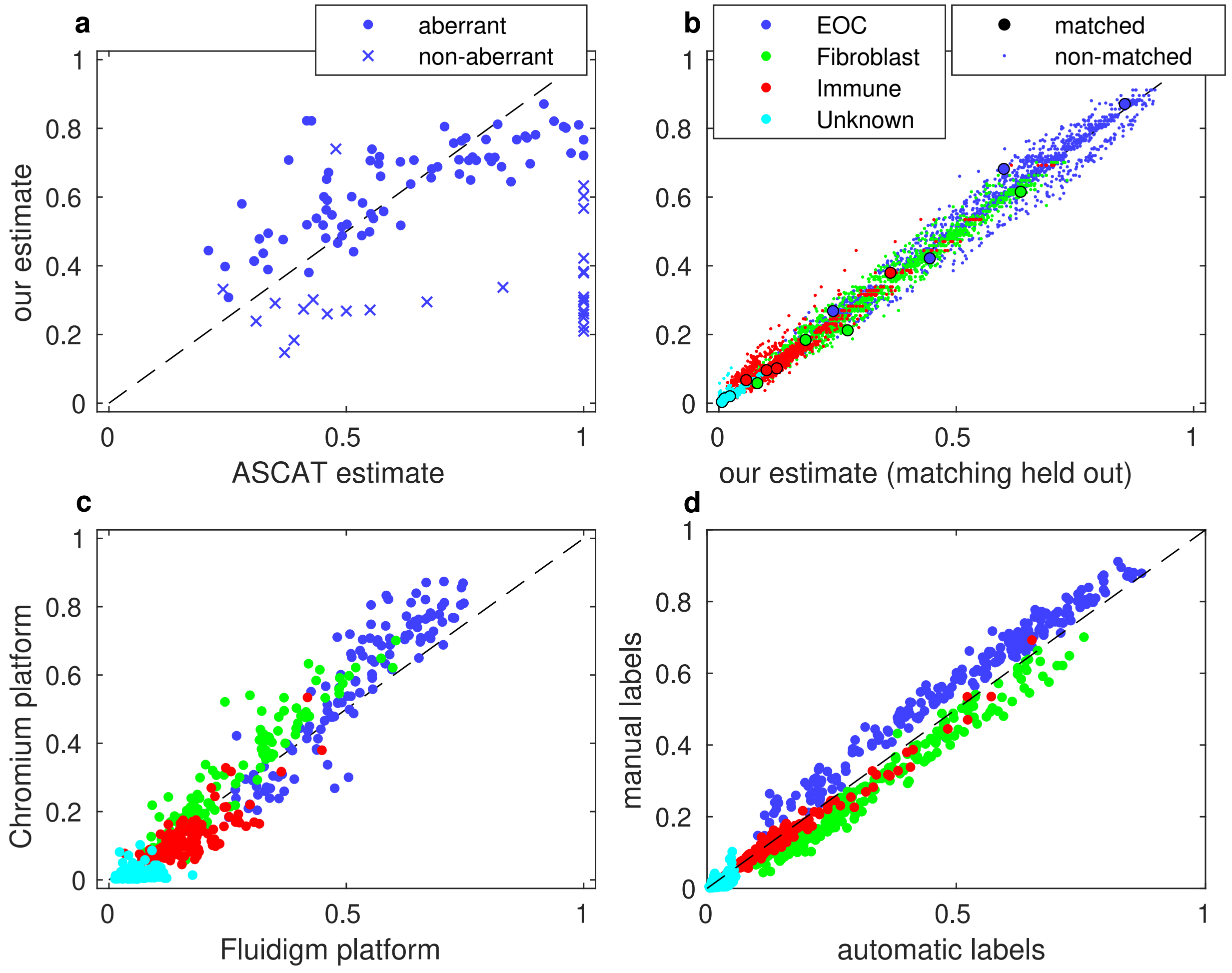


**Figure S6** Comparison of composition estimates. a) Frequency of cancer cells from whole-genome sequencing data using ASCAT [S17] and our method. The crosses represent values where the ASCAT model indicates a poor fit (reported as non-aberrant by ASCAT) [S17]. b) Frequency of each cell type using our method with matching single-cell data held out. Small dots represent all $214$ bulk samples and large dots the samples being held out. c) Frequency of each cell type using two distinct single-cell references: a $347$ cell reference from the Fluidigm platform and a $6{,}312$ cell reference from the Chromium platform. d) Composition estimated with manually-assisted labeling versus labels discovered through our clustering.

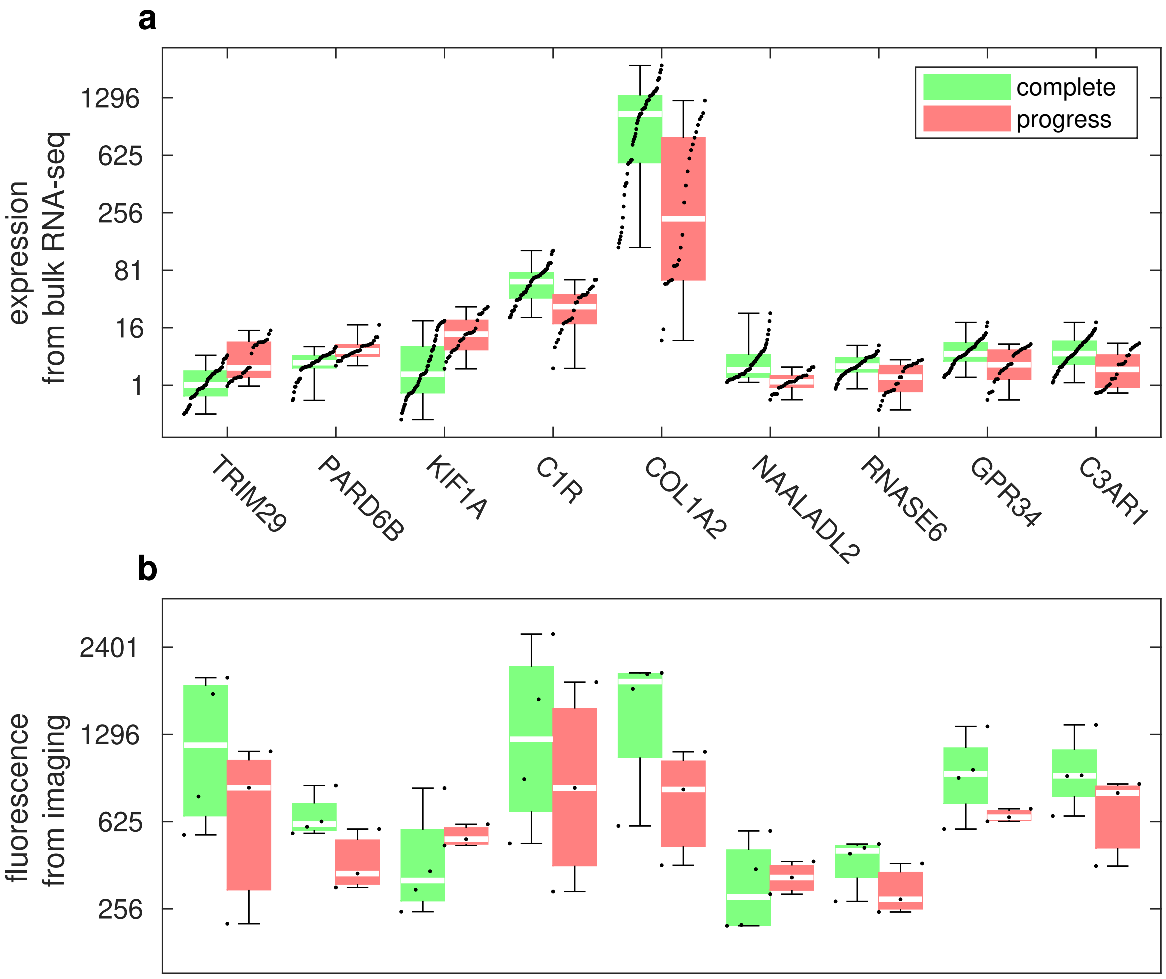


**Figure S7** Expression of the validated genes from RNA-seq and RNA *in situ* hybridization. a) Decomposed cell type specific bulk RNA expression levels, grouped by the treatment response, for all patients. b) Fluorescence intensity from RNA *in situ* hybridization, grouped by the treatment response of the patient ($4$ complete response, $3$ progressive disease). The box represents first to third quartile, white bar is median, whiskers all data, and dots are samples jittered by their rank.

$< 10^{-8}$). We also analyzed the TCGA ovarian cancer [S19] and TCGA skin cutaneous melanoma [S20] RNA sequencing dataset using solely non-matching data and the estimated sample compositions were in agreement with estimates from immunohistochemistry [S19] and genomic or methylation data [S21].

**Generalizability to other sequencing platforms.** We also compared the estimates using the 6,312 single cells from the Chromium platform versus the 347 single cell patterns from (the higher-fidelity) Fluidigm platform. Despite the smaller number of cells, the Fluidigm single-cell data features cells from 8 samples (from 8 patients), suggesting that between patient variability is similarly covered. The results are shown in bottom left panel of Figure S6 and indicate that the method works consistently across the platforms as expected. The correlation is 96.8% (p-value $< 6.2 \times 10^{-302}$) and the composition is accurately estimated even for the less frequent cell types, and the expression profiles remain consistent (rank correlation of 83.7%, p-value $< 10^{-8}$).

**Cell type specificity of expression patterns extends to pathway alterations.** Next, we analyzed whether the common pathway alterations observed in ovarian cancer [S19] tend to stem from changes of the cancer cell phenotype or if they reflect changes in the tumor microenvironment as well.

For this, we quantified the pathway activity using gene set enrichment analysis (GSEA) scores [S22] in the NCI pathway interaction database (NCI-PID) [S23] pathways from mSigDB [S24]. We computed GSEA scores using both absolute ranks (ranking the genes according to the absolute expression) and using two-tailed ranks (i.e. the median is ranked 1 and both high and low are ranked in order of increasing and decreasing expression, respectively). The former quantifies whether the genes in the pathway are expressed in general, as the expression in aspecific cell types tend to be negligible, while the latter whether the expression of the genes in the pathway is consistently aberrated, either up or down. The latter statistic tends to also indicate activity when the pathway is not employed (which itself can be an aberration, but not as interesting as being functional but perturbed; cf. Figure S9), so it is useful to inspect both. To test whether the apparent pathway activity of the bulk originate from the cancer, stromal, or immune cells we computed rank correlation between the GSEA scores from the original bulk and the decomposed, while controlling for the sample composition.

Figure S8 shows a heatmap of two-tailed GSEA scores in the NCI-PID pathways in our $214$ bulk samples. The results show that, depending on the pathway, the patterns originate from different cell types, as could be expected: the integrins and interleukins are known to be expressed in stromal and immune cell types, respectively, and our results indicate that these are mostly expressed in the corresponding cell type. Much of the signal originates from the EOC cells, which is expected, as the they are the major component in the tumor samples. Meanwhile, some patterns originate from multiple cell types, most commonly from both EOC cells and fibroblasts. The correlation with the immune signal is much lower in general due to lower immune cell content ($12.9\%$ on average), but for particular pathways and for samples with a large immune component, a strong immune signal can be detected. The large differences between the samples are explained by variations in the composition.

Next, we looked specifically at few pathways that are known to have both genomic aberra-

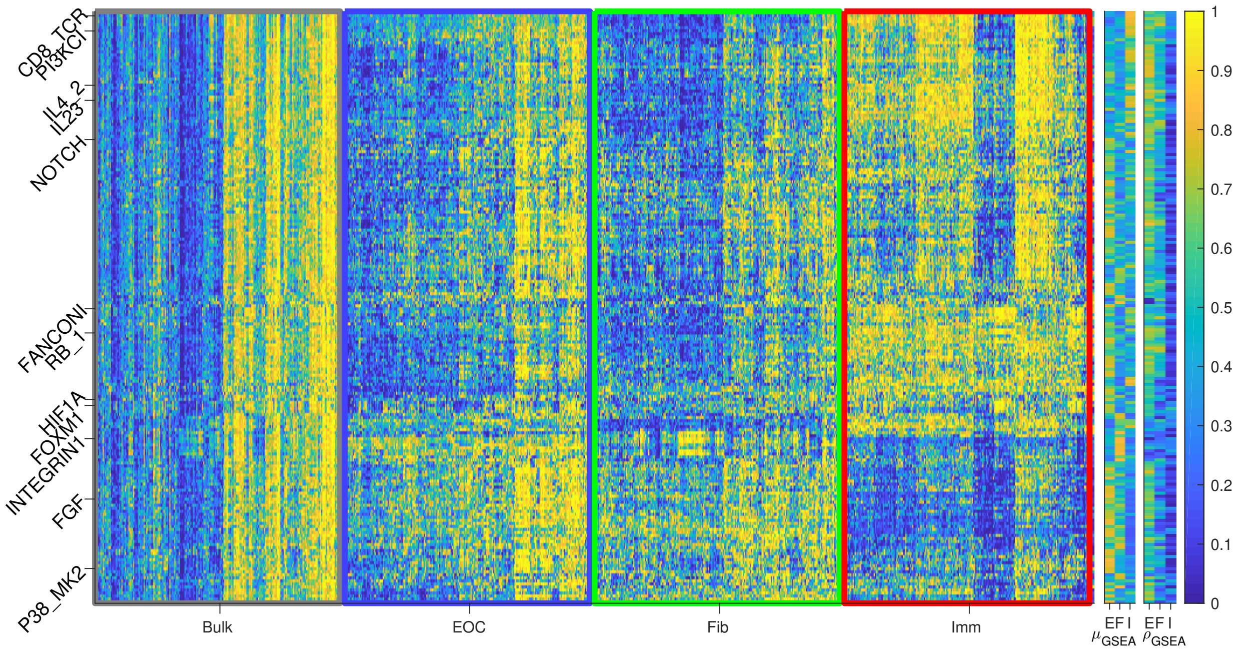


**Figure S8** Normalized two-tailed GSEA scores for the NCI-PID pathways in the HERCULES ovarian cancer samples. The plot shows the ranked two-tailed GSEA scores using the bulk and the decomposed bulk data, yellow suggesting that the pathway is either up- or downregulated with respect to the other samples. The pathways and samples are permuted according to complete linkage clustering. The flags in the $\mu_{\mathrm{GSEA}}$ indicate the average relative activity of the genes and $\rho_{\mathrm{GSEA}}$ the correlation between the bulk and the decomposed pathway activity, in each three cell type (E $\sim$ EOC, F $\sim$ fibroblasts, and I $\sim$ immune cells). Few pathways are highlighted on the y-axis.

tions and corresponding transcriptomic signal in ovarian cancer [S19]. Figure S9 shows few example pathways. Specifically, the NOTCH, FOXM1, and HIF1A pathway signals appear to originate from the cancer cells; RB1, PI3KCI (closest to PI3K/RAS pathway [S19]), and FANCONI (closest to the HR genes in [S19]) mostly from cancer and immune cells; FGF from cancer and stroma; and IL4 and IL23 mostly from immune cells. The P38_MK2, INTEGRIN1, and CD8_TCR pathways are extreme examples that are specific to a single cell type. Also in most cases, the GSEA scores are much larger in the cell type specific expression than in the bulk, indicating that the decomposition allows detecting aberrations at a finer level of detail. In the TCGA dataset, NOTCH and HIF1A signals appear to originate from the cancer cells primarily; PI3KCI from the cancer and immune cells; and FGF from cancer and stroma, validating these discoveries made using our dataset. The remaining pathways include slight differences, mostly lacking significance for the immune contribution, but we note that the TCGA samples feature even smaller immune cell fraction ($\sim 3\%$) and consequently the immune signals cannot be expected to be as pronounced.

**Tumor subtypes are confounded by sample composition.** We examined if previously discovered expression-derived tumor subtypes are affected by the sample composition. Specifically, we tested if the estimated composition varies between the subtypes, if the subtype scores correlate with a particular subtype, or if performing the subtyping only on the cancer cell profiles (i.e. disregarding the effect of stromal and immune cells) could lead to a more meaningful subtypes in terms of the patient survival.

For the ovarian cancer samples, we derived subtypes estimates using the CLOVAR method

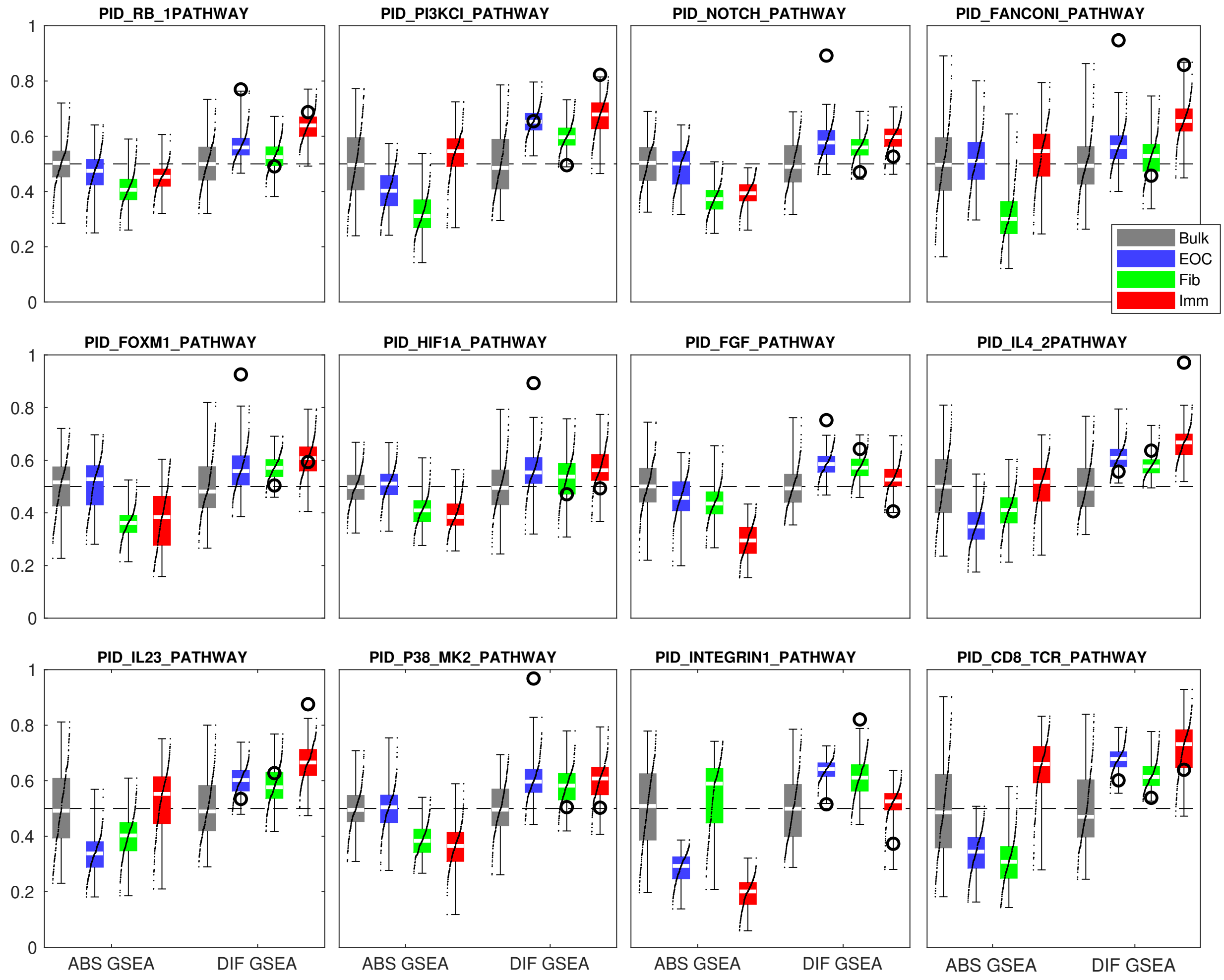


**Figure S9** Normalized one- and two-tailed GSEA scores for select pathways. The absolute (one-tailed) GSEA scores indicate whether the pathway genes are over or underexpressed in a specific samples with respect to the others, while the differential (two-tailed) GSEA scores quantify whether the genes in the pathway are consistently either up- or downregulated. The circles represent the differential GSEA score relative to the expected absolute GSEA score, indicating if the differential GSEA score is due to low or high expression in general (score of $0$) or consistent aberrations ($1$).

[S25]. In this, the samples are classified into differentiated (DIF), immunoreactive (IMR), mesenchymal (MES), or proliferative (PRO) subtype using gene set enrichment analysis [S25,S26]. The gene sets were originally discovered by clustering gene expression data from multiple ovarian cancer cohorts [S25]. Meanwhile, for the melanoma, we used the expression subtypes discovered in the TCGA melanoma study [S20]: the MITF-low (MIT), keratin (KER), and immune (IMM) subtypes. Again, we used previously reported gene lists [S20] and ssGSEA [S26] to compute null-model normalized subtype scores. For the purposes of comparison, the subtypes were also derived from the bulk data after removing components that were found to be specific to a single subtype in each setting.

Our results suggest that within the ovarian cancer subtypes, the immune subtype (IMM) is highly determined the immune cell frequency alone, while the mesenchymal subtype (MES) is greatly affect by the fibroblast frequency. Specifically, immune cells are enriched in the IMR subtype and fibroblasts in the MES subtype (p-values $< 1.2 \times 10^{-4}$ in a one-tailed rank-sum test), while varying less in the other subtypes. Moreover, a $77\%$ to $93\%$ correlation exists between the subtype score and the composition (p-values $< 10^{-8}$ in a t-test for zero correlation). Meanwhile, the frequency of cancer cells were not found to differ between the differentiated (DIF) and proliferative (PRO) subtypes (p-values $0.176$ to $0.49$), and removing the effect of fibroblasts and immune cells from the expression signal still produced consistent subtypes in these two categories, suggesting that these subtype likely reflect phenotypically different cancer cells rather than the tumor composition unlike the MES and IMR subtypes. Moreover, we found that deriving the subtypes in the absence of fibroblast and immune signals yields a better separation in the overall survival (p-value of $4.5 \times 10^{-3}$ in a log-rank test) than from the composite bulk data (p-value of $0.1483$), provided that the tumor purity is high (above median, $77\%$ for the TCGA ovarian dataset). Consistently with previous results [S25], the proliferative (PRO) subtype associated with worse survival than the differentiated (DIF), while the presence of immune cells is indicative of better survival but not significantly so (p-value $0.11$). The results were consistent between our dataset (Figure S10) and the TCGA dataset (Figure S11), but the differences in overall survival were only found significant in the TCGA dataset due to different sample sizes.

Similarly, in the skin cutaneous melanoma, we found that the immune subtype (IMM) reflects the immune cell frequency, while the MITF-low (MIT) and keratin subtypes (KER) likely reflect phenotypic differences of the cancer cells. Specifically, the fraction of immune cells is much higher in the immune subtype (p-values $< 1.8 \times 10^{-22}$, while the difference between MITF-low and keratin is barely significant (p-value $< 3.0 \times 10^{-2}$), and a strong correlation ($77\%$) exists between the immune cell frequency and the immune subtype score (p-value $< 10^{-8}$). In agreement, the results are consistent with previous findings [S20], the keratin phenotype being indicative of worse survival than the MITF-low subtype, and the differences are much stronger after removing the confounding immune component (p-value of $4.7 \times 10^{-3}$ vs $0.022$). The results are illustrated in Figure S12.

**Composition and expression of the TCGA ovarian cancer dataset.** To validate our findings, we also applied our method on bulk RNA sequencing samples from The Cancer Genome Atlas (TCGA) [S19]. We decomposed the $308$ TCGA ovarian cancer (OV) bulk-RNA samples using our (Chromium) single-cell data into three cell types: EOC, fibroblast, and immune (the cells labeled

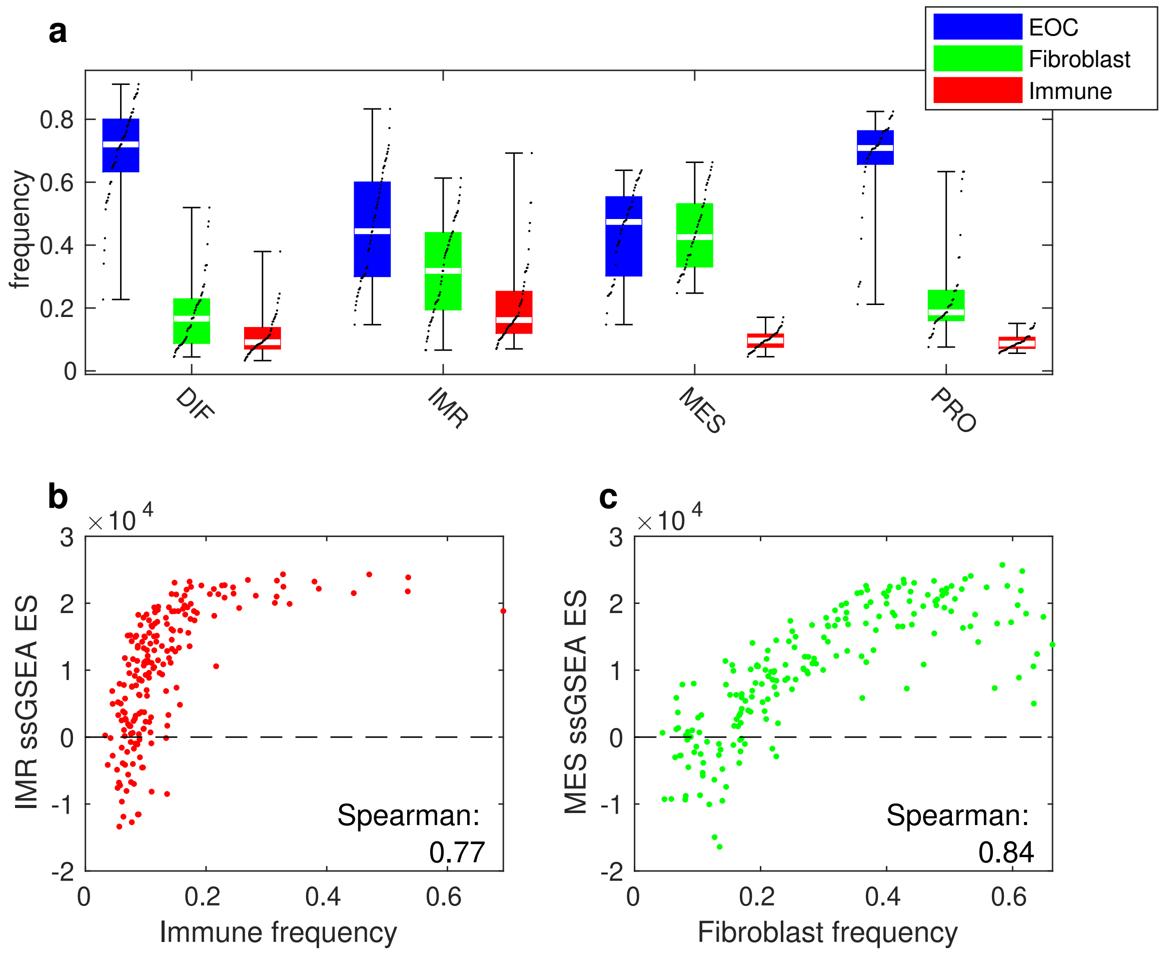


**Figure S10** Composition versus subtypes and subtype scores for the HGSOC dataset. a) Composition for each subtype: differentiated (DIF), immunoreactive (IMR), mesenchymal (MES), and proliferative (PRO). b) and c) Subtype ssGSEA score as a function of the estimated immune cell (b) and fibroblast frequency (c).

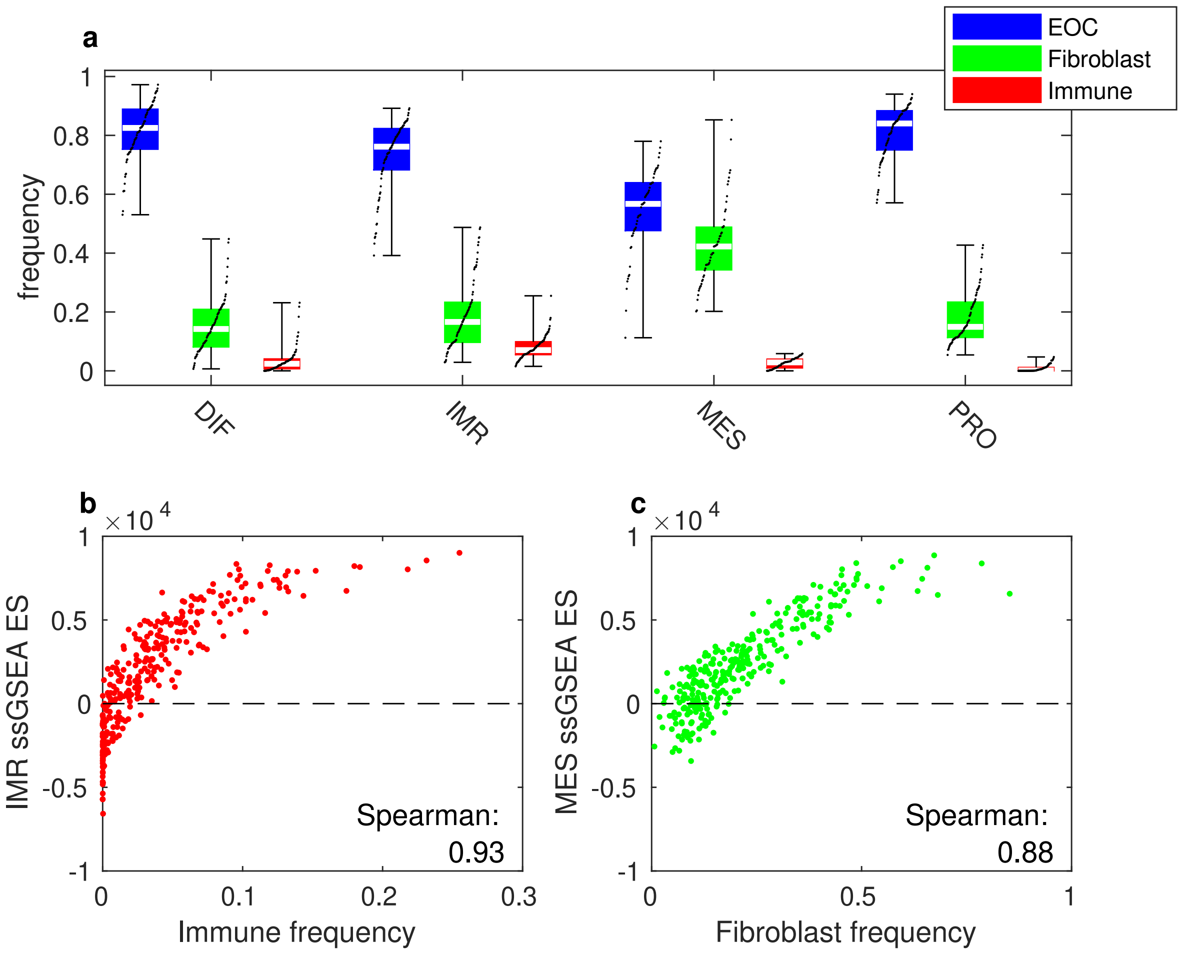


**Figure S11** Composition versus subtypes and subtype scores for the TCGA ovarian cancer dataset [S19]. a) Composition for each subtype: differentiated (DIF), immunoreactive (IMR), mesenchymal (MES), and proliferative (PRO). b) and c): subtype ssGSEA score as a function of the estimated immune cell (b) and fibroblast frequency (c).

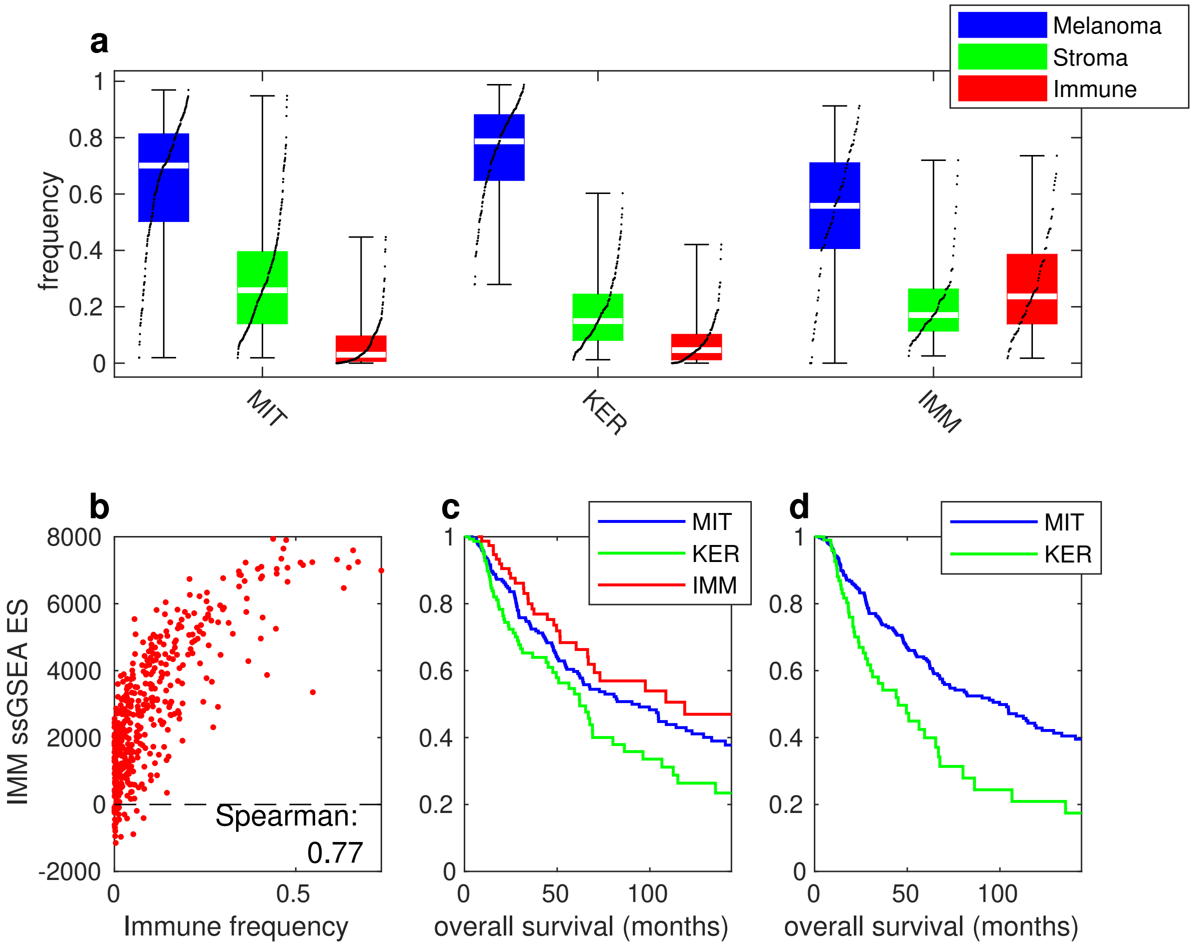


**Figure S12** Composition versus subtypes and subtype scores for the TCGA skin cutaneous melanoma dataset [S20]. a) Composition for each subtype: MITF-low (MIT), keratin (KER), and immune (IMM). b) Immune subtype ssGSEA score as a function of the estimated immune cell frequency. c) and d) Overall survival for subtypes derived from the bulk data (c) and from the bulk data after removing immune cell specific components (d).

as unknown were discarded).

First, we compared the estimated composition of these samples with those provided in the TCGA clinical data [S19] and with estimates derived from the TCGA whole-genome sequencing using ABSOLUTE [S27] by Aran et al. [S21]. Figure S13 shows the estimated compositions of the 308 bulks using the three methods. By comparing the cancer and stromal cell fractions of the TCGA clinical data to our estimates, the EOC and fibroblast frequency appears to correlate well. We find that the linear correlation between the two-methods is $86.5\%$ (p-value $< 4.3 \times 10^{-183}$ for zero linear correlation in a t-test). Meanwhile, the estimated fraction appears similar to the estimates from the genomic data using ABSOLUTE as well, the linear correalation being $61.6\%$ (p-value $< 2.5 \times 10^{-31}$). Either of these correlations is significantly better than the correlation between the TCGA clinical dat and ABSOLUTE (p-value of $< 7.4 \times 10^{-3}$ for equal correlation using Fisher's transformation and a z-test). Our estimates of EOC content are slightly lower than those from TCGA clinical data (and conversely the stromal component is lower in TCGA), but the same trend is visible in the ABSOLUTE estimates as well, which could be expected to be more accurate [S21,S27].

Next, we show that, in general, the association of gene expression with overall survival are enriched by the decomposition. Figure S14 shows the shows the pairwise comparison, and in $53.2\%$ of the cases, correlation is stronger in the decomposed (p-value $< 2.6 \times 10^{-18}$ in a binomial test). Many genes are visible in both the original bulk and in the decomposed bulk. However, some are weakly correlated in the original signal but strongly correlated after the decomposition, being

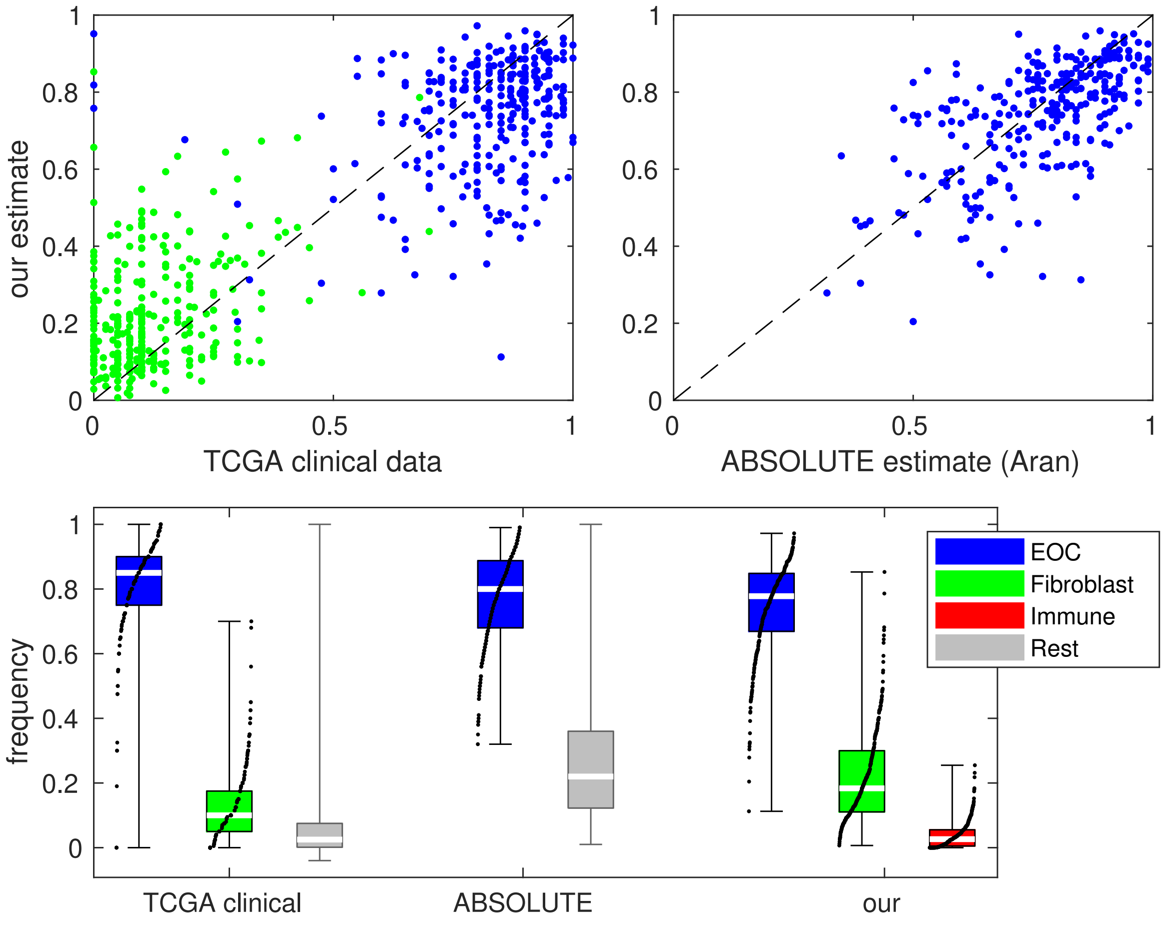


**Figure S13** Estimated composition of the TCGA ovarian cancer samples. Top left shows our estimate versus the estimated percentage of tumor and stromal cells from TCGA clinical data [S19]. Top right shows our estimate versus the estimate of tumor cells using TCGA genomic data and ABSOLUTE from Aran et al.[S21]. Bottom panel shows the distributions for each of the three methods.

prognostically favorable (e.g. COL1A2 and FCGR1A/B) or unfavorable (e.g. TEK and ERG). Some genes are also negatively correlated in the original signal, but not in the decomposed, which are due to higher-order correlations with the composition and likely represent false positive signals in the expression pattern.

Finally, we tested if there if the association with overall survival increases in the decomposed signals for the genes that were identified in our novel ovarian cancer dataset. For this, we divided the data based on the estimated expression into two ordered groups of $25\%$ of the samples each. The results are shown in Figure S15. The cancer specific genes TRIM29, PARD6B, and KIF1A feature significant association with overall survival, as suggested by our data, higher expression being less favorable. Similarly, for the stromal genes C1R and NAALADL2, a significant association can be found and the higher expression is favorable as predicted by our earlier results. The immune specific genes were not found to be significant, neither in after the decomposition or in the original bulk data. This is likely due to the lower amount of immune cells in the TCGA ovarian cancer data (estimated median of only $2.6\%$, see Figure S13) and the consequent uncertainty in the estimated immune cell profile. In general the decomposed signals tend to have stronger association as indicated by the lower p-values. We note that the C1R association is not apparent in the unprocessed data (cf. p-value of $0.2$ versus $6.6 \times 10^{-5}$ in the log-rank test).

**Composition and expression of the TCGA skin cutaneous melanoma dataset.** To verify that our method can well adapt to other cancer types, we decomposed the $474$ TCGA skin cutaneous melanoma [S20] samples using single-cell data from Tirosh et al. [S28]. The latter features a classifi-

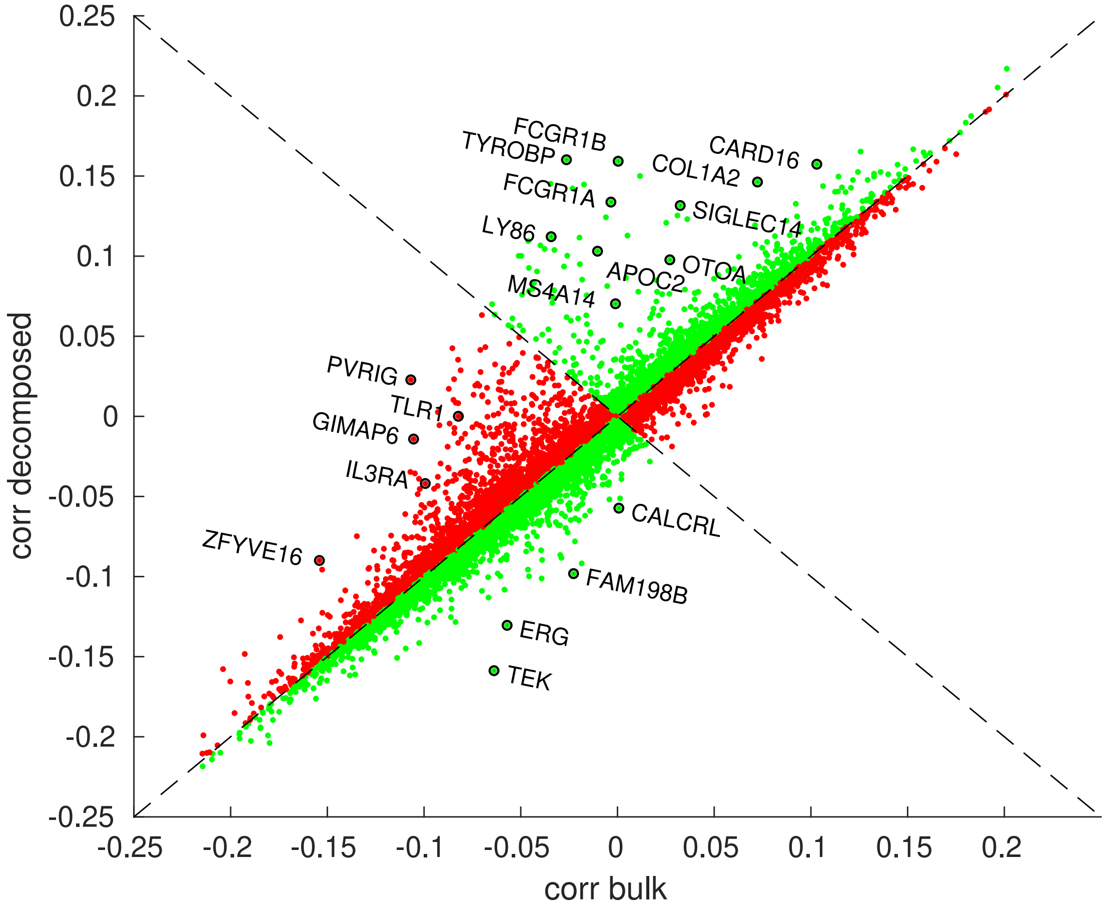


**Figure S14** Rank correlation for gene expression versus overall survival for TCGA ovarian cancer patients [S19]. Each dot represents a gene, the horizontal axis representing the correlation from undecomposed data, and the vertical axis that of the decomposed signals. Green (red) dots represent genes where the correlation is enriched (diluted) by the decomposition.

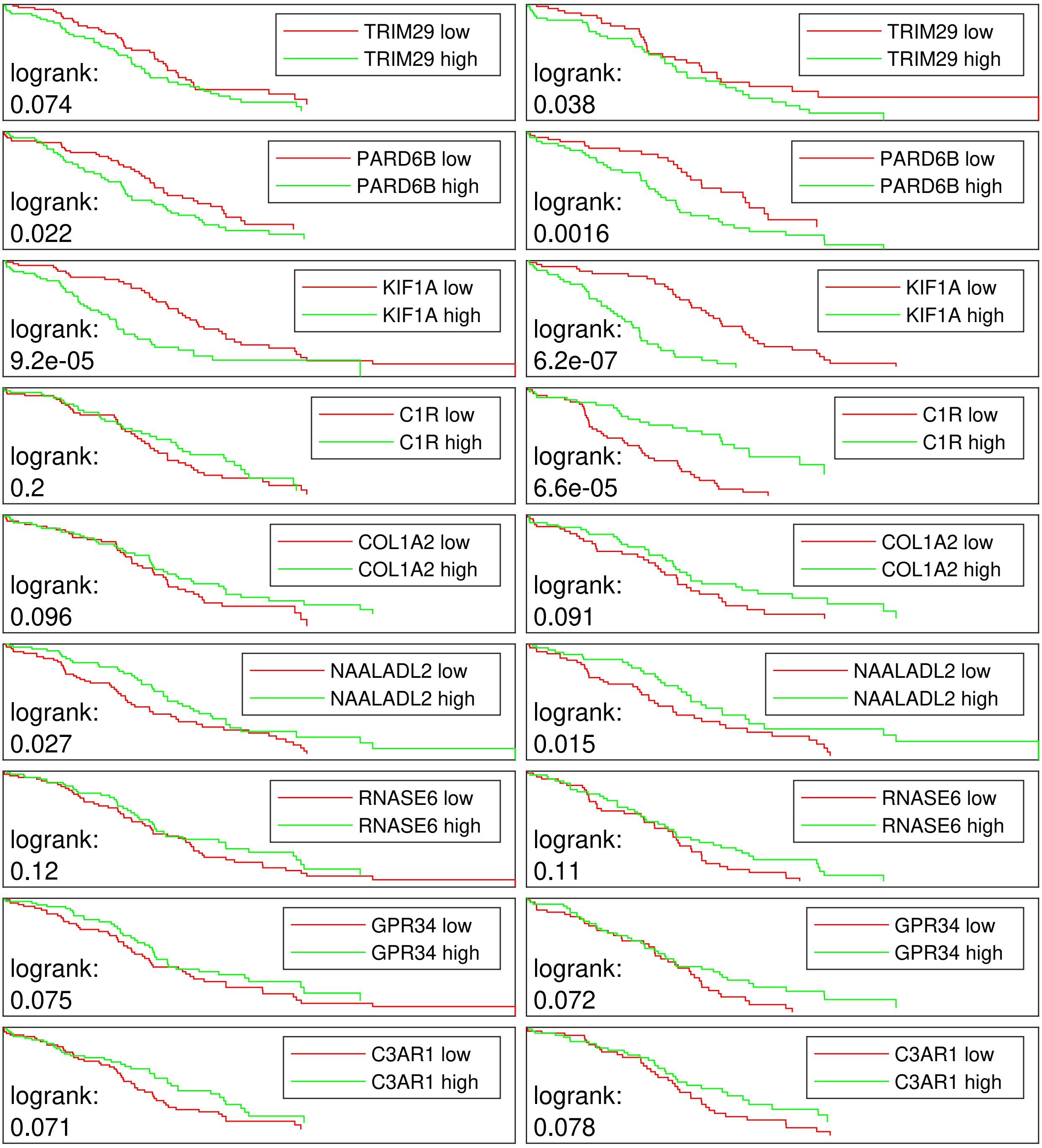


**Figure S15** Overall survival for TCGA ovarian cancer patients partitioned by gene expression (trimmed to $25\%$). Left panels: plain (undecomposed) bulk data, right panels: decomposed bulk data of the relevant cell type.

cation of the single-cells into Melanoma (1257 cells), B cells (512), T cells (2040), macrophages (119), endothelial cells (62), cancer associated fibroblasts (56), natural killers (51) (resulting in a total of 4,097 single-cells).

First, we verified that the composition is accurately estimated. As above, we compared our estimates with that of the TCGA clinical data (immunohistochemistry) and to the leukocyte unmethylation estimates (LUMP; HumanMethylome 450k array) of Aran et al. [S21]. We used the LUMP estimates as a reference, as the genomic estimates (ABSOLUTE) are not available for this cancer type and the remaining estimates in Aran et al. are not orthogonal to the immunohistochemistry and RNA sequencing platforms [S21]. The linear correlation of our estimates with those from the TCGA clinical was $71.3\%$ (p-value $< 2.1 \times 10^{-147}$ for zero linear correlation in a t-test), while the correlation with the LUMP estimates was $53.9\%$ (p-value of $< 5.2 \times 10^{-36}$). Again, both correlations are much stronger than the correlation between TCGA clinical data and the LUMP estimates (p-value of $< 6.4 \times 10^{-3}$ for equal correlation using Fisher's transformation and a z-test). The scatterplots comparing the estimates and the estimated composition of the samples are shown in Figure S16.

Next, we show that also for the melanoma dataset, the association between the expression of individual genes and the overall patient survival are generally enriched. Figure S17 shows the scatterplot of correlations in the undecomposed versus the decomposed signals. The correlations have greater magnitude in $56.6\%$ more in the decomposed versus undecomposed (p-value of $< 4.7 \times 10^{-77}$ in a binomial test). Contrary to the ovarian cancer analysis, the majority of the differ-

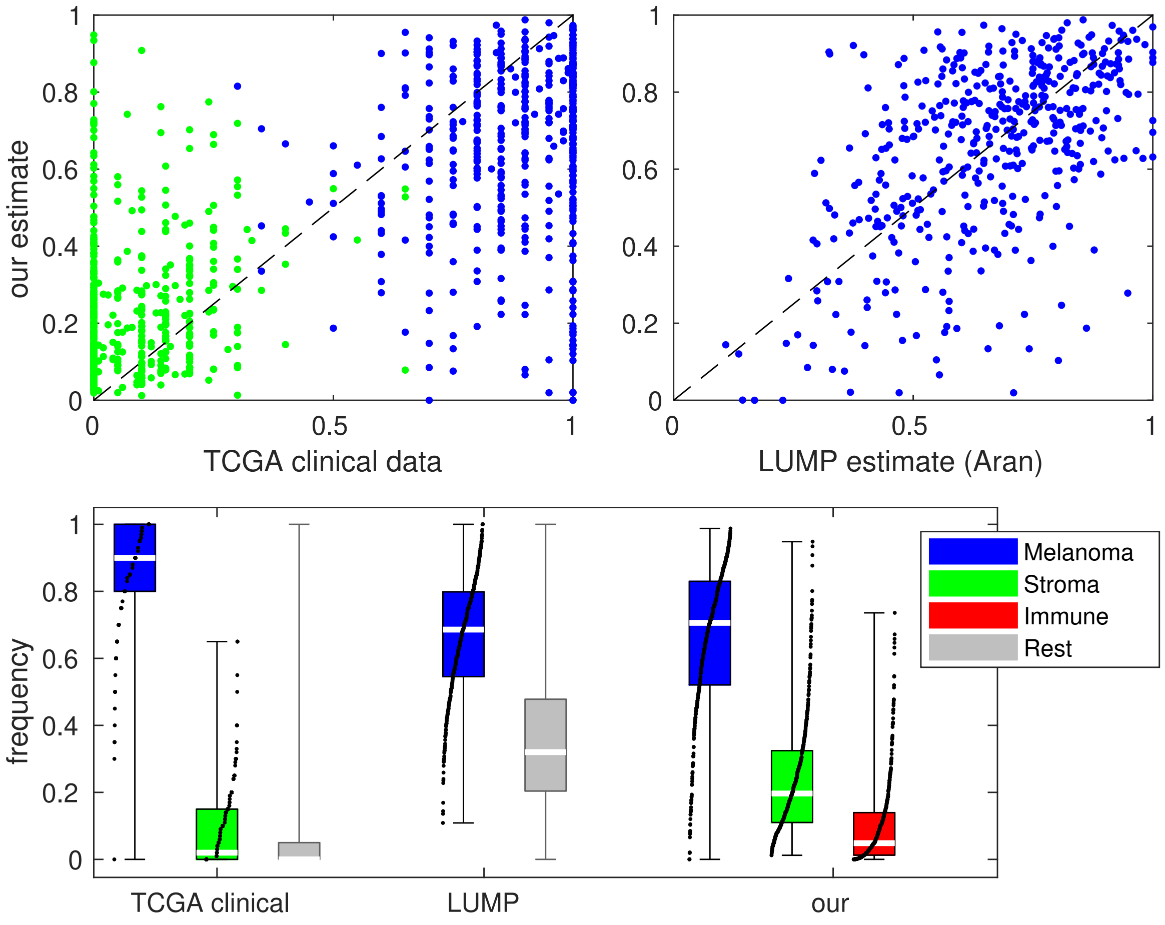


**Figure S16** Estimated composition of the TCGA skin cutaneous melanoma samples. Top left shows our estimate versus the estimated percentage of tumor and stromal cells from TCGA clinical data [S20]. Top right shows our estimate versus the estimate of tumor cells using TCGA methylome data and LUMP from Aran et al. [S21]. Bottom panel shows the distributions for each of the three methods.

ences occur in the favorable correlations, due to a much larger immune component and increased statistical power in the decomposed immune signal in single-cell data.

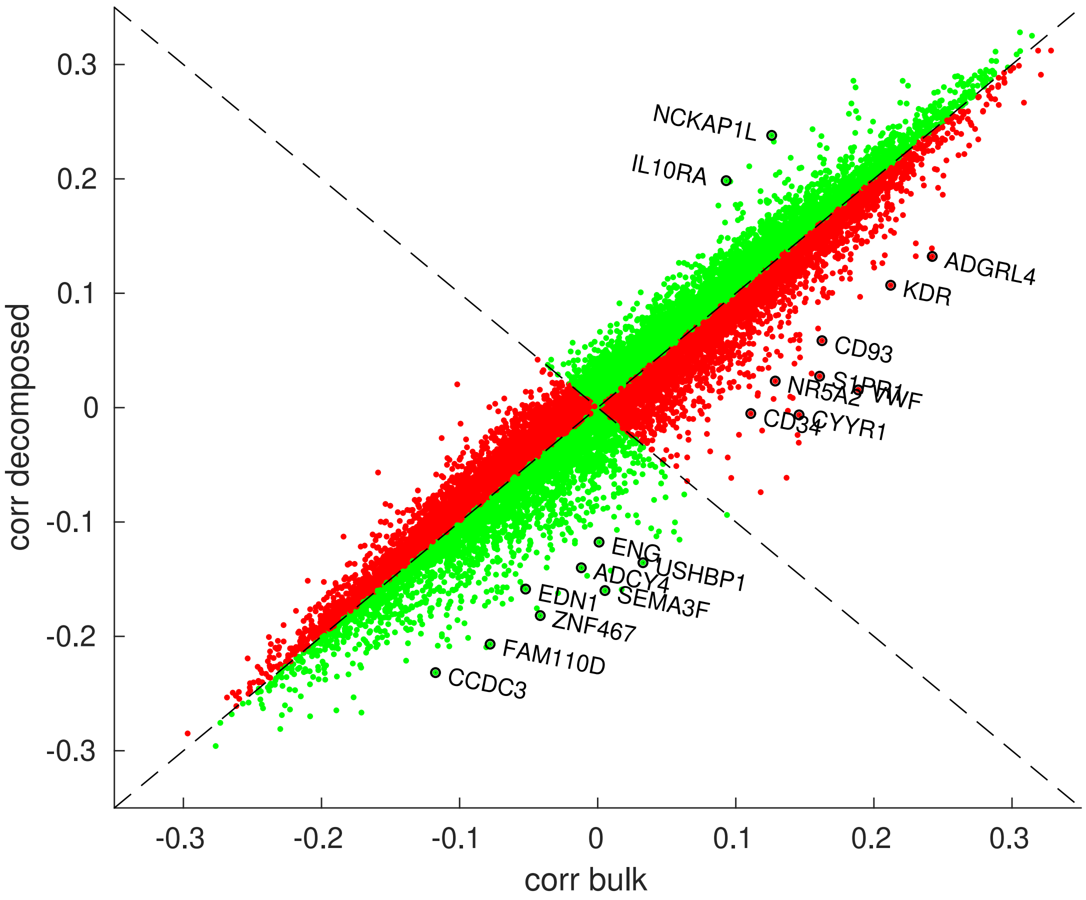


**Figure S17** Rank correlation for gene expression versus overall survival for TCGA skin cutaneous melanoma patients [S20]. Each dot represents a gene, the horizontal axis representing the correlation from undecomposed data, and the vertical axis that of the decomposed signals. Green (red) dots represent genes where the correlation is enriched (diluted) by the decomposition.

S1. Marioni, J. C., Mason, C. E., Mane, S. M., Stephens, M. & Gilad, Y. RNA-seq: an assessment of technical reproducibility and comparison with gene expression arrays. *Genome Res.* **18**, 1509–1517 (2008).

S2. McCarthy, D. J., Chen, Y. & Smyth, G. K. Differential expression analysis of multifactor RNA-Seq experiments with respect to biological variation. *Nucl. Acids Res.* **40**, 4288–4297 (2012).

S3. Raj, A. & van Oudenaarden, A. Nature, nurture, or chance: stochastic gene expression and its consequences. *Cell* **135**, 216–226 (2008).

S4. Dempster, A., Laird, N. & Rubin, D. Maximum likelihood from incomplete data via the EM algorithm. *J. Royal Stat. Soc. B* **39**, 1–38 (1977).

S5. Risso, D., Ngai, J., Speed, T. P. & Dudoit, S. Normalization of RNA-seq data using factor analysis of control genes or samples. *Nat. Biotechnol.* **32**, 896–902 (2014).

S6. Robinson, M. D. & Oshlack, A. A scaling normalization method for differential expression analysis of RNA-seq data. *Genome Biol.* **11**, R25 (2010).

S7. Zhang, Z. *et al.* SCINA: A semi-supervised subtyping algorithm of single cells and bulk samples. *Genes* **10**, 531 (2019).

S8. Zhang, A. W. *et al.* Probabilistic cell-type assignment of single-cell rna-seq for tumor microenvironment profiling. *Nat. Methods* **16**, 1007–1015 (2019).

S9. Ward, J. H., Jr. Hierarchical grouping to optimize an objective function. *J. Am. Stat. Assoc.* **58**, 236–244 (1963).

S10. Murtagh, F. A survey of recent advances in hierarchical clustering algorithms. *Comput. J.* **26**, 354–359 (1983).

S11. Yang, Y. Can the strengths of AIC and BIC be shared? a conflict between model identification and regression estimation. *Biometrika* **92**, 937–950 (2005).

S12. Bolger, A. M., Lohse, M. & Usadel, B. Trimmomatic: a flexible trimmer for Illumina sequence data. *Bioinformatics* **30**, 2114–2120 (2014).

S13. Dobin, A. *et al.* STAR: ultrafast universal RNA-seq aligner. *Bioinformatics* **29**, 15–21 (2013).

S14. Smith, T., Heger, A. & Sudbery, I. UMI-tools: modeling sequencing errors in Unique Molecular Identifiers to improve quantification accuracy. *Genome Res.* **27**, 491–499 (2017).

S15. Liao, Y., Smyth, G. K. & Shi, W. featureCounts: an efficient general purpose program for assigning sequence reads to genomic features. *Bioinformatics* **30**, 923–930 (2013).

S16. Li, H. & Durbin, R. Fast and accurate short read alignment with Burrows-Wheeler transform. *Bioinformatics* **25**, 1754–1760 (2009).

S17. Van Loo, P. *et al.* Allele-specific copy number analysis of tumors. *Proc. Natl. Acad. Sci. U.S.A.* **107**, 16910–16915 (2010).

S18. Kamentsky, L. *et al.* Improved structure, function and compatibility for CellProfiler: modular high-throughput image analysis software. *Bioinformatics* **27**, 1179–1180 (2011).

S19. The Cancer Genome Atlas Research Network. Integrated genomic analyses of ovarian carcinoma. *Nature* **474**, 609–615 (2011).

S20. The Cancer Genome Atlas Research Network. Genomic classification of cutaneous melanoma. *Cell* **161**, 1681–1696 (2015).

S21. Aran, D., Sirota, M. & Butte, A. J. Systematic pan-cancer analysis of tumour purity. *Nat. Commun.* **6**, 8971 (2015).

S22. Subramanian, A. *et al.* Gene set enrichment analysis: a knowledge-based approach for interpreting genome-wide expression profiles. *Proc. Natl. Acad. Sci. U.S.A.* **102**, 15545–15550 (2005).

S23. Schaefer, C. F. *et al.* PID: the Pathway Interaction Database. *Nucl. Acids Res.* **37**, D674–D679 (2009).

S24. Liberzon, A. *et al.* Molecular signatures database (MSigDB) 3.0. *Bioinformatics* **27**, 1739–1740 (2011).

S25. Verhaak, R. G. *et al.* Prognostically relevant gene signatures of high-grade serous ovarian carcinoma. *J. Clin. Invest.* **123**, 517–525 (2013).

S26. Barbie, D. A. *et al.* Systematic RNA interference reveals that oncogenic KRAS-driven cancers require TBK1. *Nature* **462**, 108–112 (2009).

S27. Carter, S. L. *et al.* Absolute quantification of somatic DNA alterations in human cancer. *Nat. Biotechnol.* **30**, 413–421 (2012).

S28. Tirosh, I. *et al.* Dissecting the multicellular ecosystem of metastatic melanoma by single-cell rna-seq. *Science* **352**, 189–196 (2016).
